# Paw Preference in Rats Across Tests, Strains, Sex, and Age: A PRISMA-Compliant Systematic Review and Meta-Analysis

**DOI:** 10.64898/2026.02.08.704684

**Authors:** Dipesh Pokharel, Caroline C. Swain, Dilshan H. Beligala, Madhu Vishnu Sankar Reddy Rami Reddy, Thyagarajan Subramanian

## Abstract

Paw preference in rats is widely used to study hemispheric lateralization, but many individual studies are underpowered and employ inconsistent methods, leading to conflicting reports of population-level bias. We conducted a PRISMA-compliant systematic review and meta-analysis to determine whether rats consistently display paw preference at the individual and population levels, and to evaluate the influence of behavioral test type, strain, sex, and age. Studies published between 1930 and 2025 were identified through PubMed, Google Scholar, and ScienceDirect. Data were extracted on strain, age, sex, behavioral paradigm, and paw-preference classification. Random-effects models were used to estimate pooled prevalence, with subgroup analyses for key variables. Forty studies (n = 1,609 rats) met inclusion criteria. At the individual level, 84% of rats displayed consistent paw preference (95% CI: 78–89%, p < 0.0001), demonstrating robust individual-level lateralization. However, population-level analyses showed no universal directional bias, right paw use occurred in 48% of rats (95% CI: 43–54%) and left paw use in 39% (95% CI: 34–44%). Ambidextrous classification thresholds were standardized across studies to ensure comparability. Subgroup analyses indicated modest strain- and test-dependent effects, with Sprague Dawley rats tending toward balanced paw use, while other strains showed slight rightward bias. Skilled-reaching tasks produced slightly stronger asymmetry than the Collins test. Sex- and age-related differences were subtle and inconsistent. Overall, rats exhibit reliable individual-level paw preference without species-wide directional asymmetry, distinguishing them from humans. Standardized testing protocols and balanced cohort designs will enhance reproducibility and translational relevance in lateralization research.

## 1. Introduction

Lateralization of brain function, first described in humans, is now recognized as an evolutionary conserved feature across species, from invertebrates to vertebrates.^1^ Hemispheric specialization enhances neural efficiency by reducing redundancy and enabling parallel processing of cognitive and motor functions.^2^ One of the most accessible behavioral readouts of this specialization is limb or paw preference, a simple and quantifiable marker of motor asymmetry. Such asymmetries are observed across primates, birds, reptiles, canines, rodents, and even insects, underscoring lateralization as a fundamental principle of nervous system organization.^3–10^ In humans, handedness is the most prominent manifestation, with ∼90% of individuals right-handed, a population-wide asymmetry linked to left-hemispheric dominance for language and fine motor control.^11–14^ In rodents, however, population-level findings are inconsistent. While many studies report robust and stable paw preferences at the individual level, reports of directional bias across groups vary widely, with some indicating rightward preference, others leftward, and many finding no asymmetry.^15–21^

These discrepancies likely reflect small sample sizes, strain-specific factors, and variability in behavioral assays.^16, 17, 22^ Beyond ethological interest, rodent paw preference has important translational implications. It has been linked to asymmetries in dopaminergic signaling within the nigrostriatal pathway, vulnerability to stress-induced neuroplasticity, and immune function differences.^23–30^ Such associations are particularly relevant for modeling neurological disorders with asymmetric symptoms before progressing bilaterally.^31–37^ Paw preference may therefore serve as a behavioral marker of hemispheric lateralization with direct relevance to neurodegenerative disease models.^38–42^

Despite nearly a century of research, several key questions remain unresolved. Do rats consistently exhibit stable paw preferences at both the individual and population levels? To what extent do strain, sex, age, and testing paradigm influence the strength or direction of lateralization? Prior meta-analyses have pooled results across species, thereby obscuring species-specific patterns.^16, 17, 43^ A focused synthesis of rat-only data is therefore essential to clarify whether population-level asymmetry exists and to identify sources of experimental variability.

Also, addressing these gaps is essential for improving reproducibility in behavioral neuroscience and refining rodent models in translational research.^16, 44^ To address this, we conducted a PRISMA-compliant systematic review and meta-analysis of rat paw preference across nearly a century of research (1930–2025). Our aims were to: (1) determine whether rats consistently display paw preference at individual and population levels; and (2) evaluate the extent to which strain, sex, age, and behavioral test paradigm shape outcomes. Through this synthesis, we clarify the reliability of paw preference as a marker of hemispheric specialization, identify sources of heterogeneity, and assess its translational relevance for modeling asymmetry-related neurological disorders.

## 2. Methods

### 2.1 Protocol

The protocol for this systematic review and meta-analysis was performed according to the PRISMA guideline and registered in PROSPERO (Registration #CRD42025621247).^45, 46^

### 2.2 Eligibility criteria

We included studies of all rat strains, ages, and sexes in which paw preference was assessed using validated behavioral tests. Non-rat studies were excluded. Review articles, preprints, and dissertations were included if they contained extractable primary data. Only English-language publications were eligible. Unpublished or inaccessible studies were excluded.

### 2.3 Information sources and Search strategy

Electronic searches were conducted in PubMed, Google Scholar, and ScienceDirect. The initial search was performed in December 2023 and updated in October 2024.

The PubMed search syntax was as follows:

(“Rats” [MeSH] OR rat OR rats) AND (Functional Laterality) “pawedness” [tiab] OR “paw preference“[tiab] OR “limb preference“[tiab] OR handedness[tiab] OR lateralization[tiab] OR “limb dominance“[tiab] OR “dominant paw“[tiab] OR “preferred paw“[tiab] OR “limb laterality“[tiab] “Limb asymmetry“[tiab] OR “motor asymmetry“[tiab] OR “motor laterality“[tiab]

### 2.4 Selection process

The study selection process is illustrated in a PRISMA 2020 flow diagram. (Figure 1) Two independent reviewers trained in behavioral neuroscience, under the supervision of a movement disorders specialist (T.S.), completed the initial screening. Duplicates and ineligible studies were removed based on titles, abstracts, and methods to determine paw preference. Each study was then screened independently by two reviewers to determine eligibility based on specific inclusion and exclusion criteria to determine the final list of studies to be included. For any disagreements, a third screener determined eligibility. Figure 1 outlines the flow of study selection from databases.

**Figure 1:**
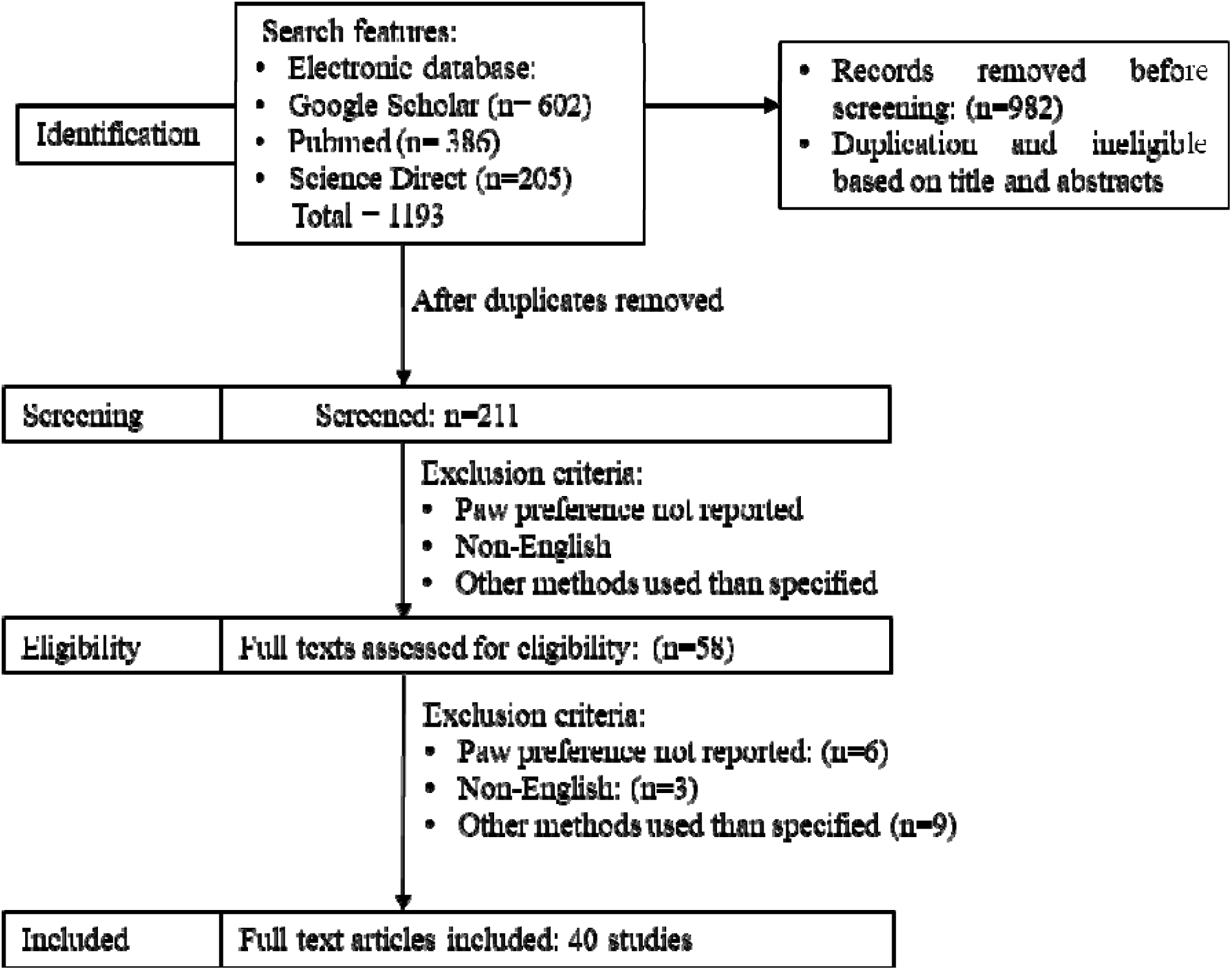
PRISMA 2020 flow diagram for new systematic reviews which included searches of databases and registers only. Source : Page MJ, et al. BMJ 2021;372:n71. doi: 10.1136/bmj.n71.

### 2.5 Data Extraction

Two reviewers independently extracted study characteristics and outcome data into a standardized excel sheet (Table 1). Extracted variables included study title, authors, publication year, rat strain, age, sex, behavioral test type, and paw preference outcomes (right, left, or ambidextrous). When data were available only in graphical form, values were digitized using graphreader.com/v2.^47^ For studies reporting multiple experimental groups, all groups were included. Effect measures were derived from reported or extracted outcomes. If the same cohort was tested longitudinally, data from the earliest time point were extracted to minimize the influence of training effects.^48–52^ Discrepancies between reviewers were resolved by consensus, with arbitration by a third reviewer when necessary.

**Table 1:** Summary of included studies with paw preference distributions by strain, sex, age, and test type.

| Study | Strains | Sex | Age | Test | Left paw preference | Right paw preference | Ambidextrous | Total |
| --- | --- | --- | --- | --- | --- | --- | --- | --- |
| Agestam & Cahusac_2007<sup>59</sup> | Wistar rat | Male | Adult | Platform reaching | 0.0 | 6.0 | 4.0 | 10.0 |
| Allred et al._2006<sup>60</sup> | Sprague<br>Dawley rat | Male | Adult | Vermicelli handling | 16.0 | 13.0 | 66.0 | 95.0 |
| Larson_1981<sup>61</sup> | Sprague<br>Dawley rat | Female | Young | Collins | 13.0 | 11.0 | 4.0 | 28.0 |
| de Souza Pagnussat et al._2009<sup>62</sup> | Wistar rat | Male | Adult | Staircase | 19.0 | 31.0 | 2.0 | 52.0 |
| Elamis et al._2003<sup>63</sup> | Wistar rat | Female | Adult | Collins | 7.0 | 56.0 | 5.0 | 68.0 |
| Guyen et al._2003<sup>64</sup> | Wistar rat | Mixed | Adult | Collins | 144.0 | 39.0 | 15.0 | 198.0 |
| Hernandez-Mesa et al._1985<sup>65</sup> | Hooded | Male | Adult | Collins | 6.0 | 7.0 | 2.0 | 15.0 |
| Kurtlu et al._2012<sup>66</sup> | Wistar | Female | Adult | Modified food-reaching | 14.0 | 31.0 | 5.0 | 50.0 |
| Martin & Webster_1974<sup>49</sup> | Unknown | Male | Young | Collins modified | 28.0 | 18.0 | 1.0 | 47.0 |
| Miklyeva & Bures_1991<sup>67</sup> | Long-Evans<br>hooded rat | Male | Adult | Collins | 27.0 | 20.0 | 6.0 | 53.0 |
| Miklyeva et al._1991<sup>50</sup> | Hooded | Mixed | Young | Collins | 76.0 | 60.0 | 38.0 | 174.0 |
| Milsen_1937<sup>68</sup> | Unknown | Mixed | Adult | Circular-cage test | 0.0 | 4.0 | 1.0 | 5.0 |
| Pence_2002<sup>69</sup> | Sprague<br>Dawley rat | Mixed | Adult | Collins | 22.0 | 80.0 | 12.0 | 114.0 |
| Peterson strain 1_1931<sup>70</sup> | Unknown | Mixed | Adult | Skilled reaching | 2.0 | 3.0 | 0.0 | 5.0 |
| Peterson strain 2_1931<sup>70</sup> | Hooded | Male | Adult | Skilled reaching | 1.0 | 1.0 | 0.0 | 2.0 |
| Stashkevich & Kulikov_2001<sup>51</sup> | Wistar rat | Mixed | Young | Collins | 24.0 | 23.0 | 14.0 | 61.0 |
| Sullivan et al._2012<sup>71</sup> | Long-Evans rat | Mixed | Adult | Lateral paw | 24.0 | 27.0 | 2.0 | 53.0 |
| Tsai & Maurer_1930<sup>72</sup> | Unknown | Mixed | Young | Circular cage | 33.0 | 55.0 | 18.0 | 106.0 |
| Volnova & Kurzina_2013<sup>73</sup> | Wistar rats | Mixed | Adults | Collins | 9.0 | 7.0 | 3.0 | 19.0 |
| Whishaw et al._1986<sup>52</sup> | Long Evans rat | Male | Adult | Skilled reaching | 25.0 | 18.0 | 21.0 | 64.0 |
| Wu et al._2010<sup>74</sup> | Wistar rat | Male | Young | Collins | 12.0 | 39.0 | 9.0 | 60.0 |
| Ambeskovic et al._2017<sup>75</sup> | Long-Evans rat | Male | Adult | Collins | 12.0 | 33.0 | 0.0 | 45.0 |
| Cunha et al._2017<sup>76</sup> | Sprague Dawley rat | Male | Adult | Pawedness trait | 11.0 | 13.0 | 0.0 | 24.0 |
| Dunnett et al._1987<sup>77</sup> | Sprague Dawley rat | Female | Young | Skilled reaching | 10.0 | 9.0 | 0.0 | 19.0 |
| Ellens et al._2016<sup>78</sup> | Sprague Dawley rat | Male | Adult | Skilled reaching | 7.0 | 11.0 | 0.0 | 18.0 |
| Evenden & Robbins_1984<sup>79</sup> | Lister Hooded rat | Male | Adult | Skilled reaching | 13.0 | 11.0 | 0.0 | 24.0 |
| O'Bryan et al. strain 1_2011<sup>80</sup> | Wistar rat | Male | Adult | Staircase Test | 50.0 | 49.0 | 0.0 | 99.0 |
| O'Bryan et al. strain 2_2011<sup>80</sup> | Wistar rat | Mixed | Adult | Collins | 76.0 | 89.0 | 0.0 | 165.0 |
| O'Bryan et al. strain 3_2011<sup>80</sup> | Long Evans | Male | Adult | Lateral paw | 28.0 | 32.0 | 0.0 | 60.0 |
| Peterson_1938<sup>81</sup> | Unknown | Male | Adult | Lever pressing | 8.0 | 9.0 | 0.0 | 17.0 |
| Peterson_1951<sup>82</sup> | White rat | Male | Adult | Collins | 21.0 | 14.0 | 0.0 | 35.0 |
| Schwartz et al._1987<sup>83</sup> | Wistar | Male | Adult | Lever pressing | 3.0 | 4.0 | 0.0 | 7.0 |
| Siegfried & Bures_1980<sup>84</sup> | Wistar rat | Male | Adult | Collins | 47.0 | 38.0 | 45.0 | 130.0 |
| Stashkevich & Kulikov_2010<sup>85</sup> | Wistar | Male | Adult | Collins | 11.0 | 16.0 | 0.0 | 27.0 |
| Stashkevich & Kulikov_2007<sup>86</sup> | Wistar | Male | Adult | Collins | 38.0 | 48.0 | 0.0 | 86.0 |
| Vyazovskiy & Tobler_2008<sup>87</sup> | Sprague Dawley rat | Male | Adult | Collins | 5.0 | 5.0 | 0.0 | 10.0 |
| Ecevitoglu, Alev, et al._2018<sup>23</sup> | Sprague Dawley rat | Mixed | Adult | Collins modified | 9.0 | 9.0 | 6.0 | 24.0 |
| Groneberg Elena, et al._2025<sup>88</sup> | Lister Hooded rat | Mixed | Adult | Collins | 14.0 | 22.0 | 0.0 | 36.0 |
| Pokharel Dipesh, et al._2025<sup>22</sup> | Sprague Dawley rat | Mixed | Mixed | Collins | 68.0 | 75.0 | 30.0 | 173.0 |
| Parmiani, Pierantonio, et al._2021<sup>89</sup> | Wistar rat | Male | Adult | Skilled reaching | 6.0 | 2.0 | 2.0 | 10.0 |

**Notes:** *Mixed groups included studies in which sex or age was not separated (e.g., both males and females, or mixed-age cohorts). These were retained in the pooled prevalence analyses to preserve statistical power but were excluded from sex- or age-specific subgroup comparisons. Unknown groups where strain, sex, or age were not reported were included in the overall meta-analysis to ensure comprehensive representation of historical data but were excluded from stratified analyses requiring those variables (e.g., strain-, sex-, or age-based comparisons). This approach-maintained comparability across studies while minimizing data loss and bias introduced by inconsistent reporting.*

### 2.6 Ethical Statement

This study did not involve live animals and was based solely on published data; therefore, institutional ethical approval was not required.

### 2.7 Study risk of bias assessment

Results of risk of bias assessment are summarized in Figure 2 (traffic-light plot) and detailed in Table 2. Risk of bias was assessed using SYRCLE’s risk of bias tool for animal studies, adapted from the Cochrane tool.^53^ This tool evaluates 10 domains of bias, each of which is rated as *low risk (Yes)*, *unclear risk*, or *high risk (No)*. Types of bias assessed are selection, performance, detection, attrition, reporting, and other bias. Each study was assessed by two independent reviewers (two medical students and one postdoctoral fellow). Discrepancies were resolved by a third reviewer.

**Figure 2.**
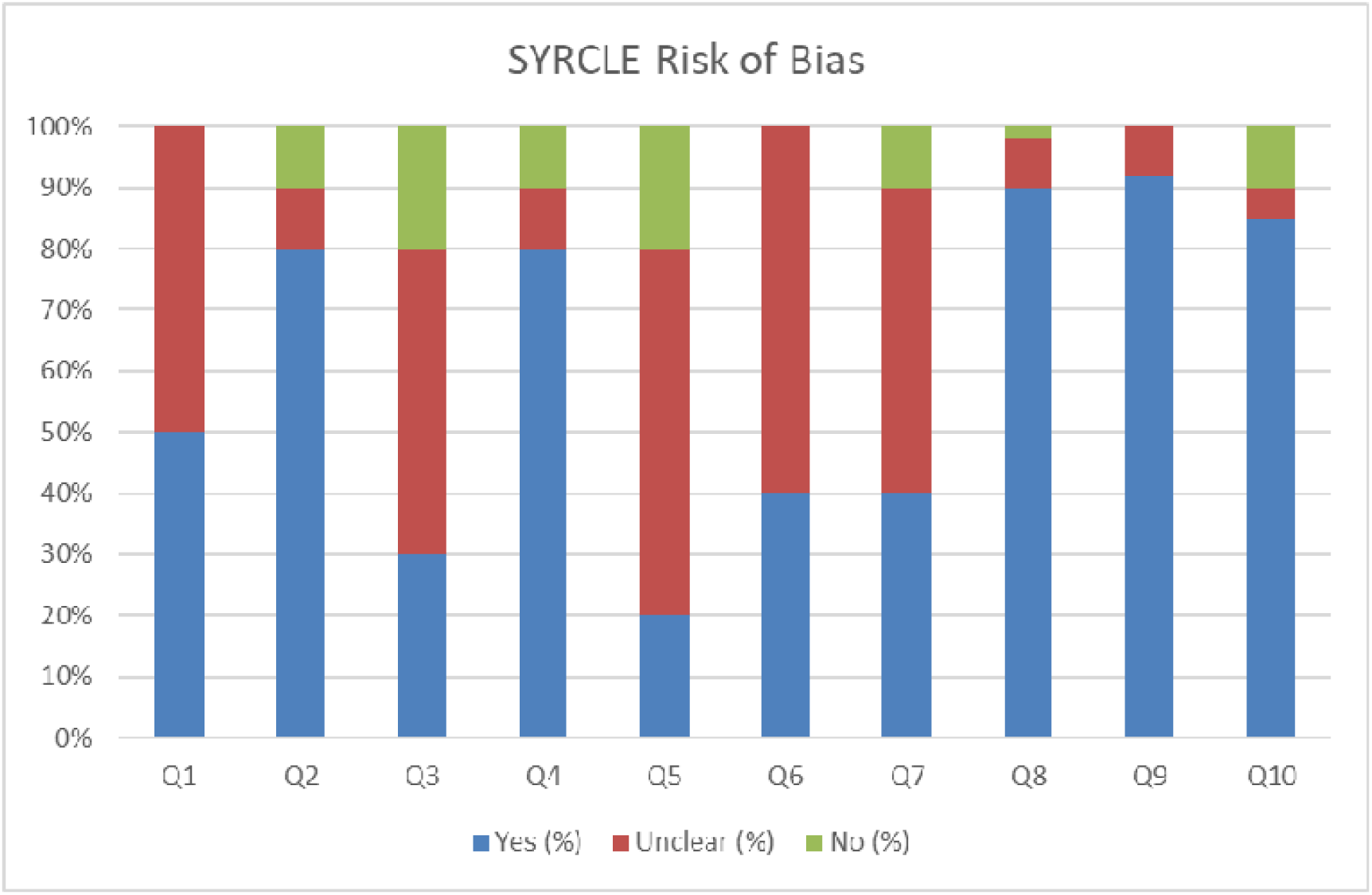
SYRCLE Risk of Bias Summary. Proportions of studies rated as low, unclear, or high risk of bias across the 10 SYRCLE domains. Green = low risk, yellow = unclear risk, red = high risk across ten methodological domains for included studies.

**Table 2.**
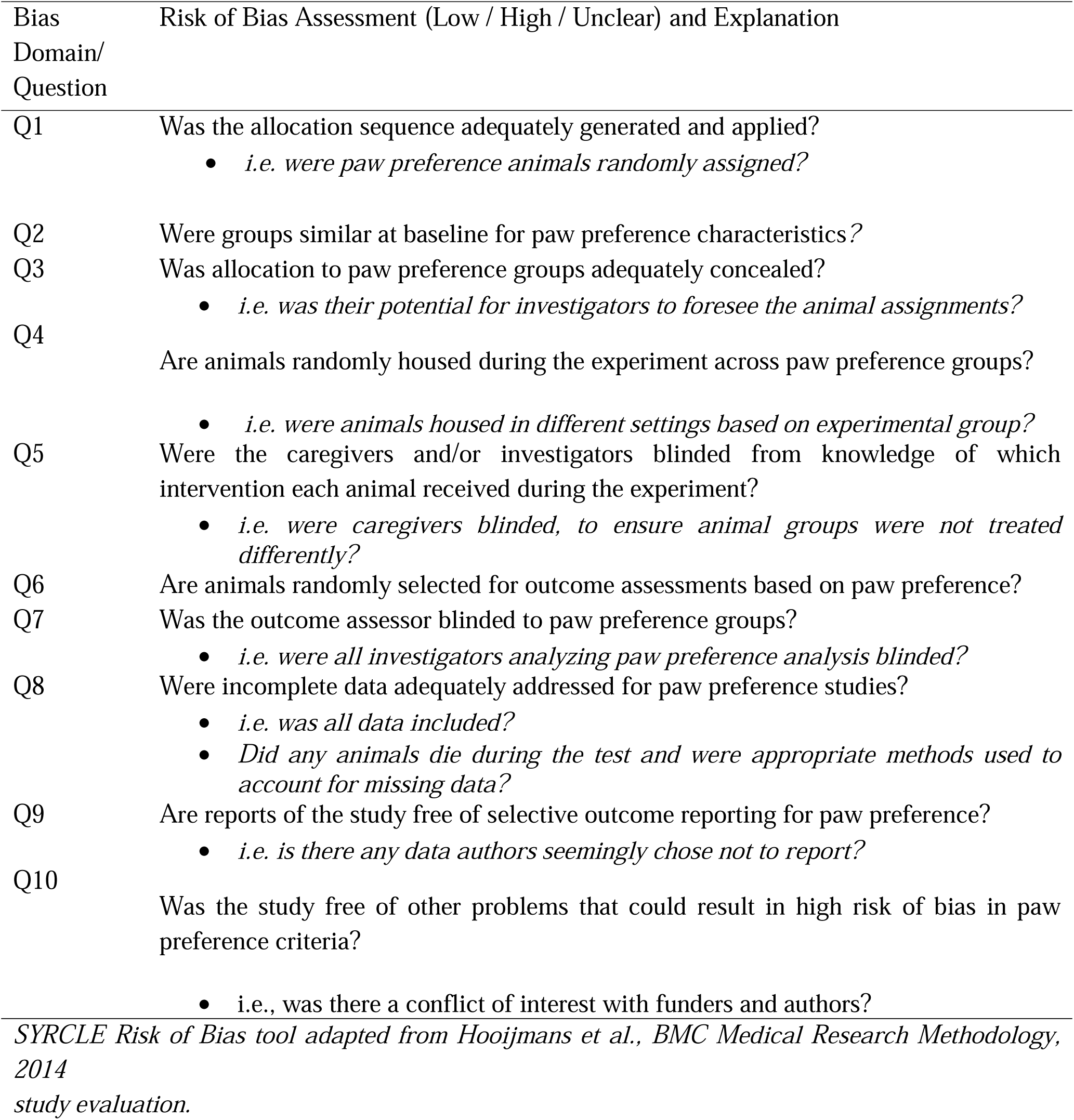
SYRCLE Risk of Bias domains and paw-preference specific operational criteria used for.

| Bias Domain/<br>Question | Risk of Bias Assessment (Low / High / Unclear) and Explanation |
| --- | --- |
| Q1 | Was the allocation sequence adequately generated and applied?<br>*i.e. were paw preference animals randomly assigned?* |
| Q2 | Were groups similar at baseline for paw preference characteristics? |
| Q3 | Was allocation to paw preference groups adequately concealed?<br>*i.e. was their potential for investigators to foresee the animal assignments?* |
| Q4 | Are animals randomly housed during the experiment across paw preference groups?<br>*i.e. were animals housed in different settings based on experimental group?* |
| Q5 | Were the caregivers and/or investigators blinded from knowledge of which intervention each animal received during the experiment?<br>*i.e. were caregivers blinded, to ensure animal groups were not treated differently?* |
| Q6 | Are animals randomly selected for outcome assessments based on paw preference? |
| Q7 | Was the outcome assessor blinded to paw preference groups?<br>*i.e. were all investigators analyzing paw preference analysis blinded?* |
| Q8 | Were incomplete data adequately addressed for paw preference studies?<br>*i.e. was all data included?*<br>*Did any animals die during the test and were appropriate methods used to account for missing data?* |
| Q9 | Are reports of the study free of selective outcome reporting for paw preference?<br>*i.e. is there any data authors seemingly chose not to report?* |
| Q10 | Was the study free of other problems that could result in high risk of bias in paw preference criteria?<br>*i.e., was there a conflict of interest with funders and authors?* |

*SYRCLE Risk of Bias tool adapted from Hooijmans et al., BMC Medical Research Methodology, 2014 study evaluation.*

### 2.8 Synthesis methods

#### 2.8.1 Meta-Analysis Model

All analyses employed random-effects models to account for between-study variability. Pooled prevalence estimates with corresponding 95% confidence intervals (CIs) were computed, and comparisons against chance (0.5) were evaluated using Z-tests.

We first examined hemispheric asymmetry at the individual level, including only studies that categorized paw preference into three groups: *left*, *right*, and *ambidextrous*. Event rates were calculated for lateralized (left + right) versus non-lateralized (ambidextrous) animals.

Next, hemispheric asymmetry was analyzed at the population level, incorporating all eligible studies. Two separate meta-analyses were performed: Event rates were used as effect measures. Right-pawedness vs. non–right-pawedness (left + ambidextrous) and Left-pawedness vs. non–left-pawedness (right + ambidextrous).

**Note:** The total pooled sample size across all analyses (n = 2,288 rats) was greater than the number of unique animals (n = 1,609) because separate right- and left-paw event datasets were analyzed independently at the population-level. This approach maintains comparability with prior laterality meta-analyses and ensures accurate estimation of directional bias.

#### 2.8.2 Subgroup Comparisons

Planned subgroup analyses examined:

- Test paradigm: Collins Test vs. other validated tests
- Strain: Sprague Dawley rat vs. other strains
- Sex: Male and female rats
- Age: Young and adult/old rats

#### 2.8.3 Stratified Analyses

Additional stratified analyses compared paw preference within sex × strain groups (e.g., male Sprague Dawley rat vs. male other strains; female Sprague Dawley rat vs. female other strains). Both right and left paw preference proportions were examined separately.

#### 2.8.4 Test Paradigm Comparisons

Collins test was directly compared against other validated paw preference tests across all strains to evaluate methodological influences on lateralization.

#### 2.8.5 Handling Missing Data

When necessary, missing numerical values (e.g., n, mean, or Standard error of the mean) (SEM) were imputed using mean values derived from studies with comparable characteristics. Standard deviations (SDs) were calculated from reported SEM and sample size (n). When only a range of n was provided, the smallest reported value was used to maintain a conservative estimate. If only body weight was reported, chronological age was extrapolated using published strain-specific age/weight reference charts.^54–56^ Conversely, if only age ranges were provided, the median value was used for analysis.

#### 2.8.6 Age Classification

To ensure uniformity across studies that variably reported age or body weight, rats were categorized as young or adult based on standardized developmental criteria.

- Young: 150–240 g; 5–8 weeks old (< 1.8 months)
- Adult: 260–800 g; 9–48 weeks old (> 2 months)

This classification was applied uniformly during subgroup analyses to minimize bias arising from inconsistent age or weight reporting. By harmonizing developmental categories across datasets, age-related effects could be more accurate compared to studies using differing reporting conventions.

#### 2.8.7 Harmonization of Ambidextrous Cut-offs

To ensure methodological consistency, ambidextrous classification thresholds were harmonized based on conventions established in prior meta-analytic studies of rodent paw preference.^16, 17^ We adopted their three-category framework (left, ambidextrous, right) and aligned each study’s original criteria such as right paw events (RPE) ranges or asymmetry coefficients with this standardized schema. When thresholds were explicitly stated in the primary source (e.g., RPE ≤ 21 = left; 22–28 = ambidextrous; ≥ 29 = right), they were retained verbatim. For studies that reported alternative metrics (e.g., asymmetry coefficients, Z- or Wilcoxon-based thresholds, or task-specific rules), equivalent left/ambidextrous/right categories were derived from the authors’ descriptions. This harmonization ensured that each study contributed comparable categorical data to the pooled analysis while preserving original decision criteria. We then updated the evidence base by incorporating additional rat studies published between 2020 and 2025, using identical inclusion/exclusion criteria and search strategies described above. Newly identified studies were screened in duplicate and cross-verified for data accuracy. After harmonization, meta-analytic computations were restricted to *Rattus norvegicus* to generate species-specific pooled estimates.

### 2.9 Tabulation and graphical methods

Forest plots were generated to display pooled estimates for each comparison (strain, sex, and test type). Forest plots (Figures 3–23) summarize pooled right, left, and ambidextrous paw preference proportions, stratified by strain, sex, and test paradigm. Summary pooled estimates are presented in Supplementary Table 1.

**Figure 3.**
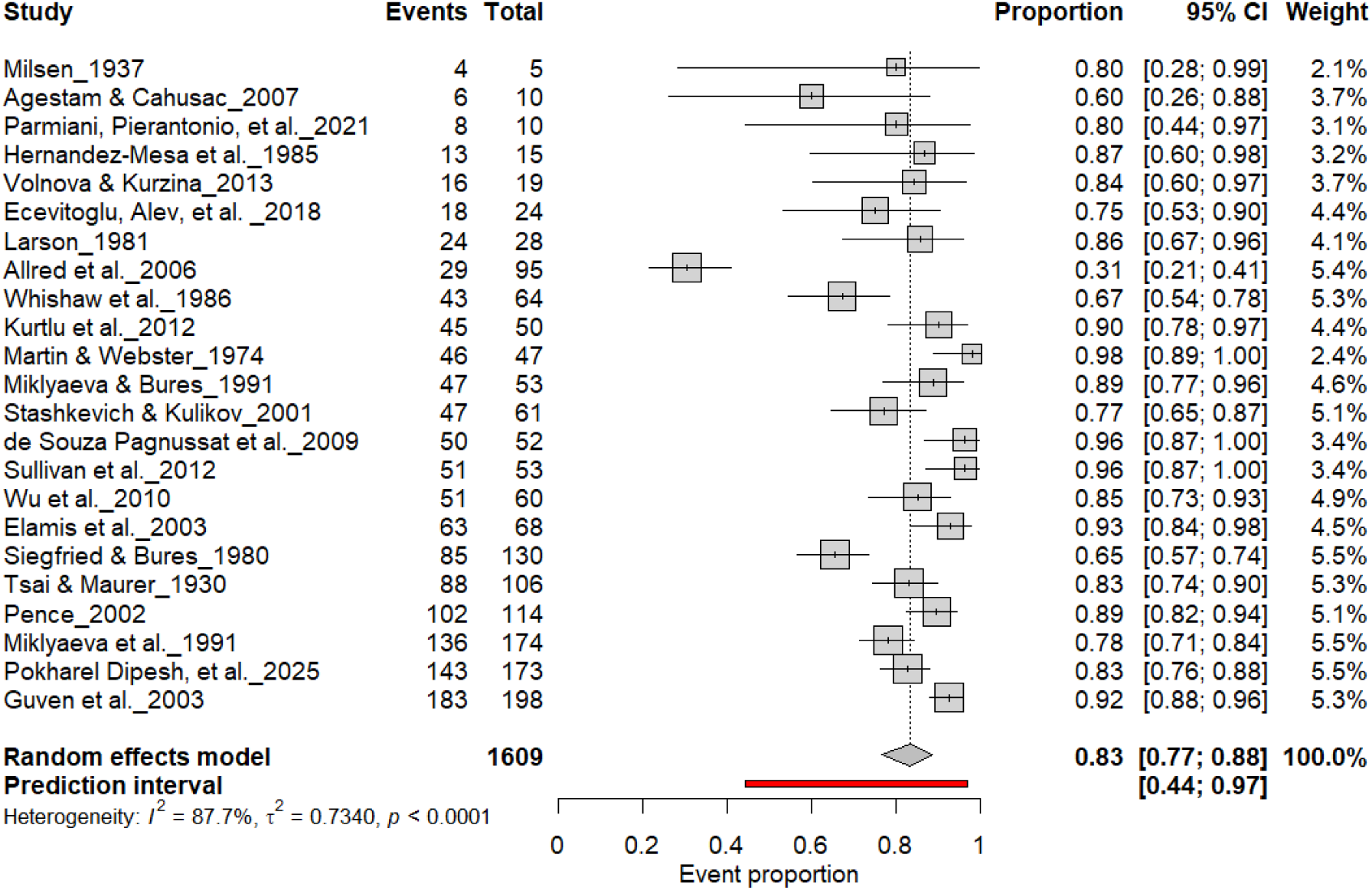
Forest plot of 24 studies (n = 1,609 rats) showing the proportion of animals exhibiting consistent paw preference. Each square denotes an individual study estimate (size reflects weight), with horizontal lines representing 95% confidence intervals. The diamond indicates the pooled estimate from a random-effects model, demonstrating that 83% of rats exhibit stable individual-level lateralization (95% CI: 0.77–0.88, p < 0.0001) despite substantial heterogeneity (I² = 87.7%).

**Figure 4.**
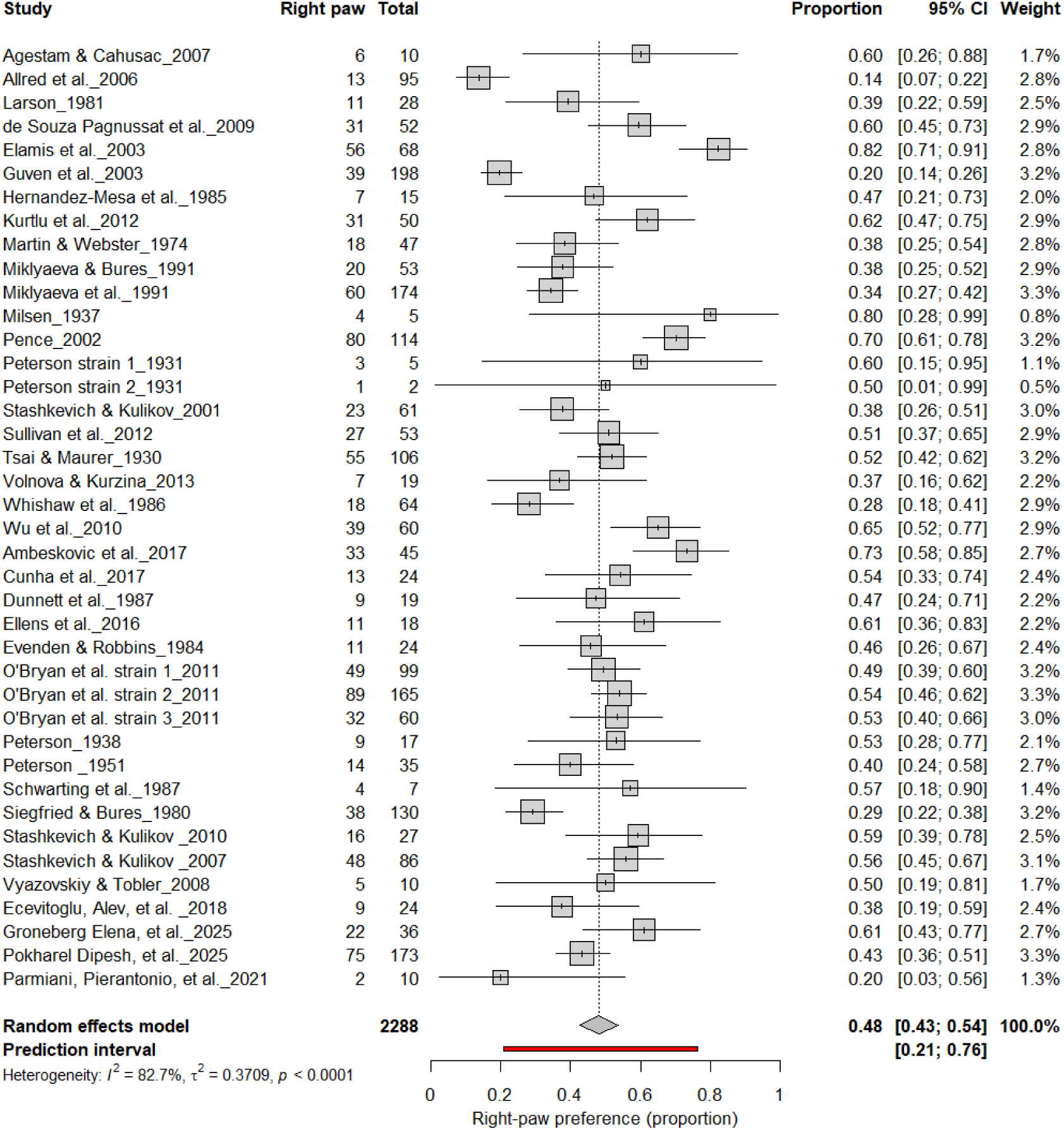
Forest plot of 40 studies (n = 2,288 rats) showing the proportion of animals classified as right pawed. Each square represents the study-specific estimate (size proportional to study weight), and horizontal lines indicate 95% confidence intervals. The diamond represents the pooled estimate derived from a random-effects model, indicating that 48% of rats were right-pawed (95% CI: 0.43–0.54, p < 0.0001). Substantial between-study heterogeneity was observed (I² = 82.7%), with no evidence of a population-level rightward bias.

**Figure 5.**
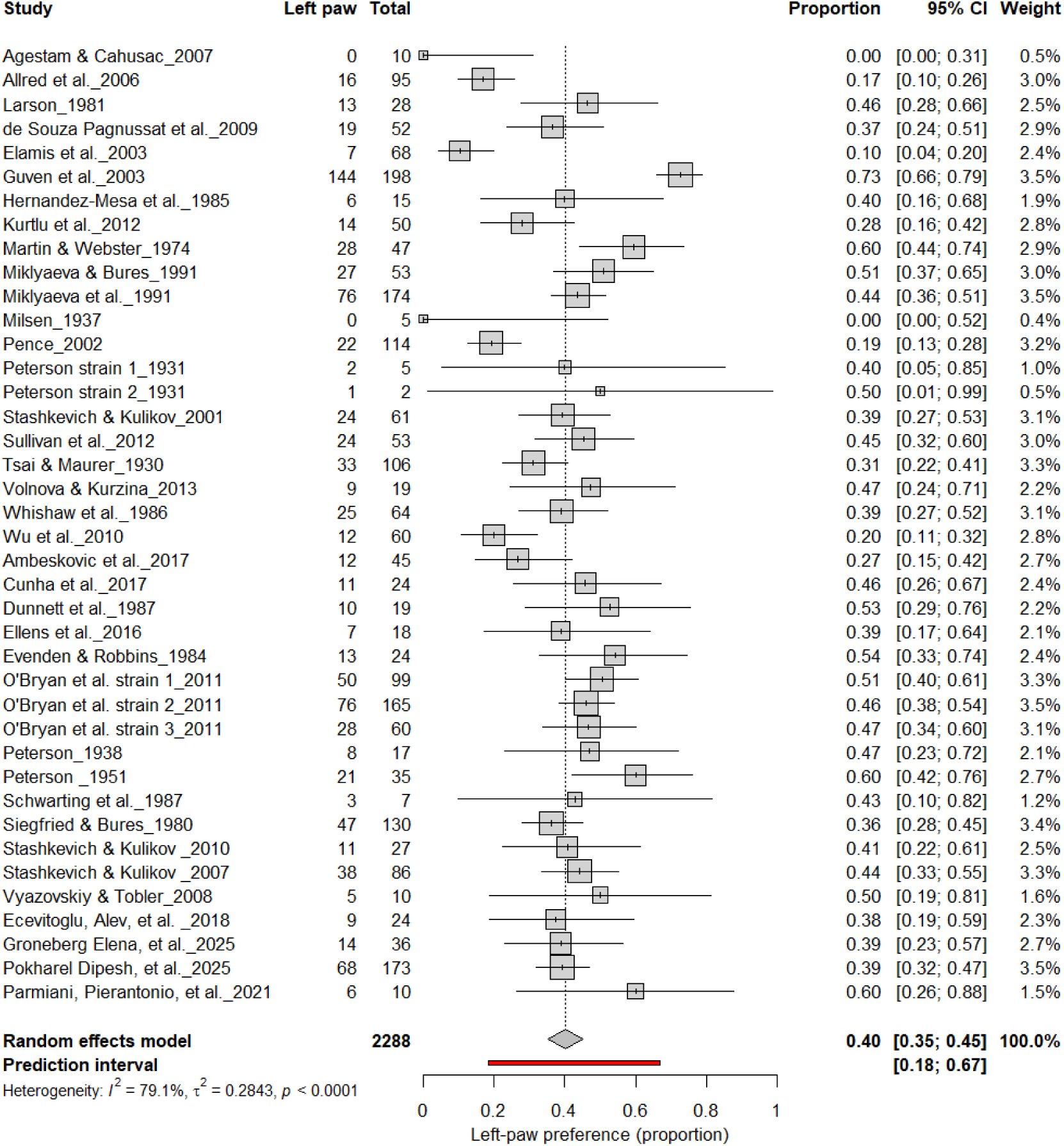
Forest plot of 40 studies (n = 2,288 rats) showing the proportion of animals classified as left pawed. Each square represents the study-specific estimate (size proportional to study weight), and horizontal lines denote 95% confidence intervals. The diamond indicates the pooled estimate from a random-effects model, showing that 40% of rats were left-pawed (95% CI: 0.35–0.45, p < 0.0001). Substantial between-study heterogeneity was observed (I² = 79.1%), with no evidence of a population-level leftward bias.

**Figure 6.**
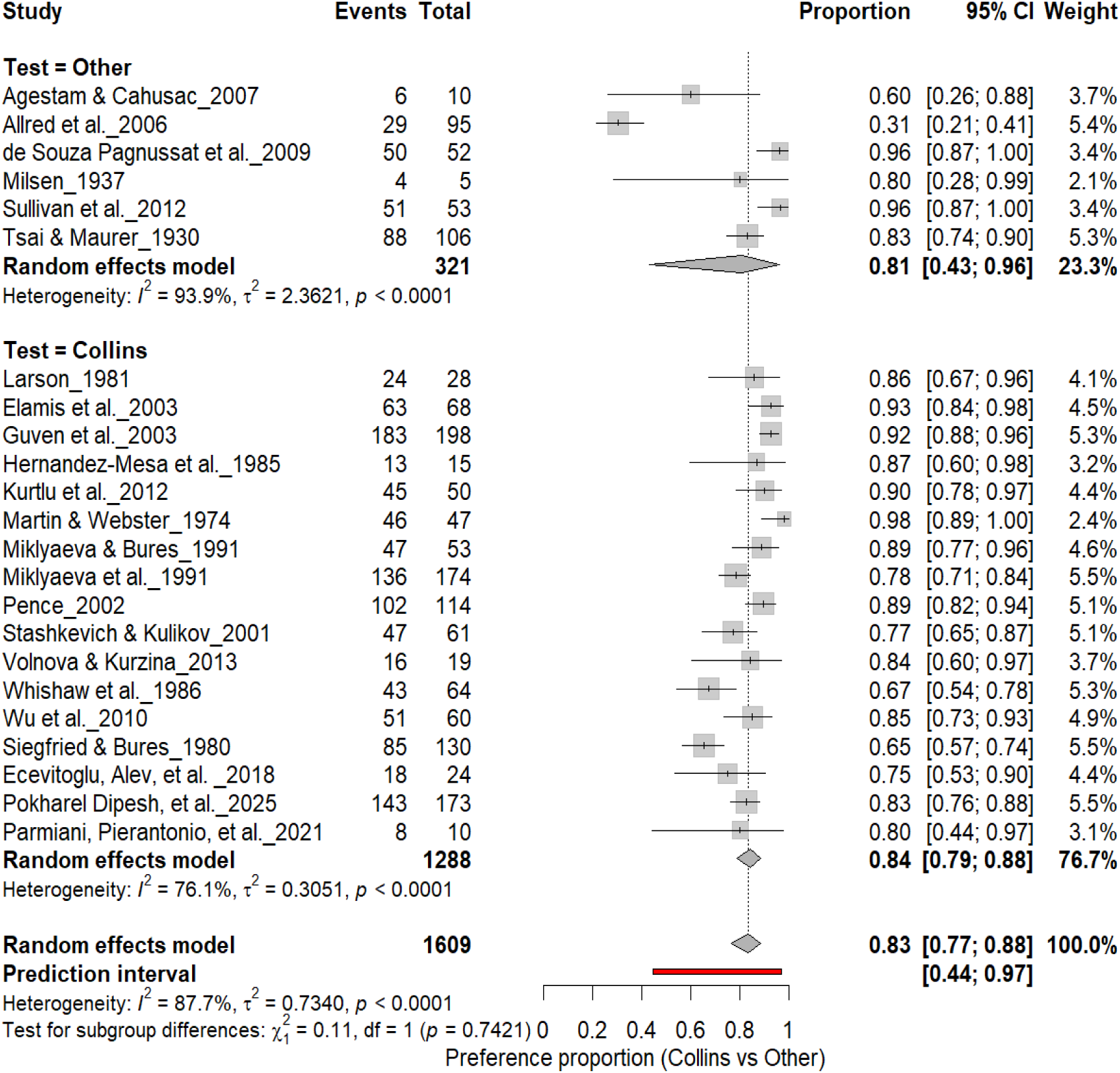
Forest plot comparing the Collins test with other behavioral paradigms for assessing individual-level paw preference. Each square represents the study-specific estimate (size proportional to study weight), with horizontal lines indicating 95% confidence intervals. Both subgroups showed a high prevalence of lateralization (Collins: 83%; Other tests: 81%), with no significant subgroup difference (χ² = 0.11, df = 1, p = 0.74). Substantial heterogeneity was observed across studies (I² = 87.7%).

**Figure 7.**
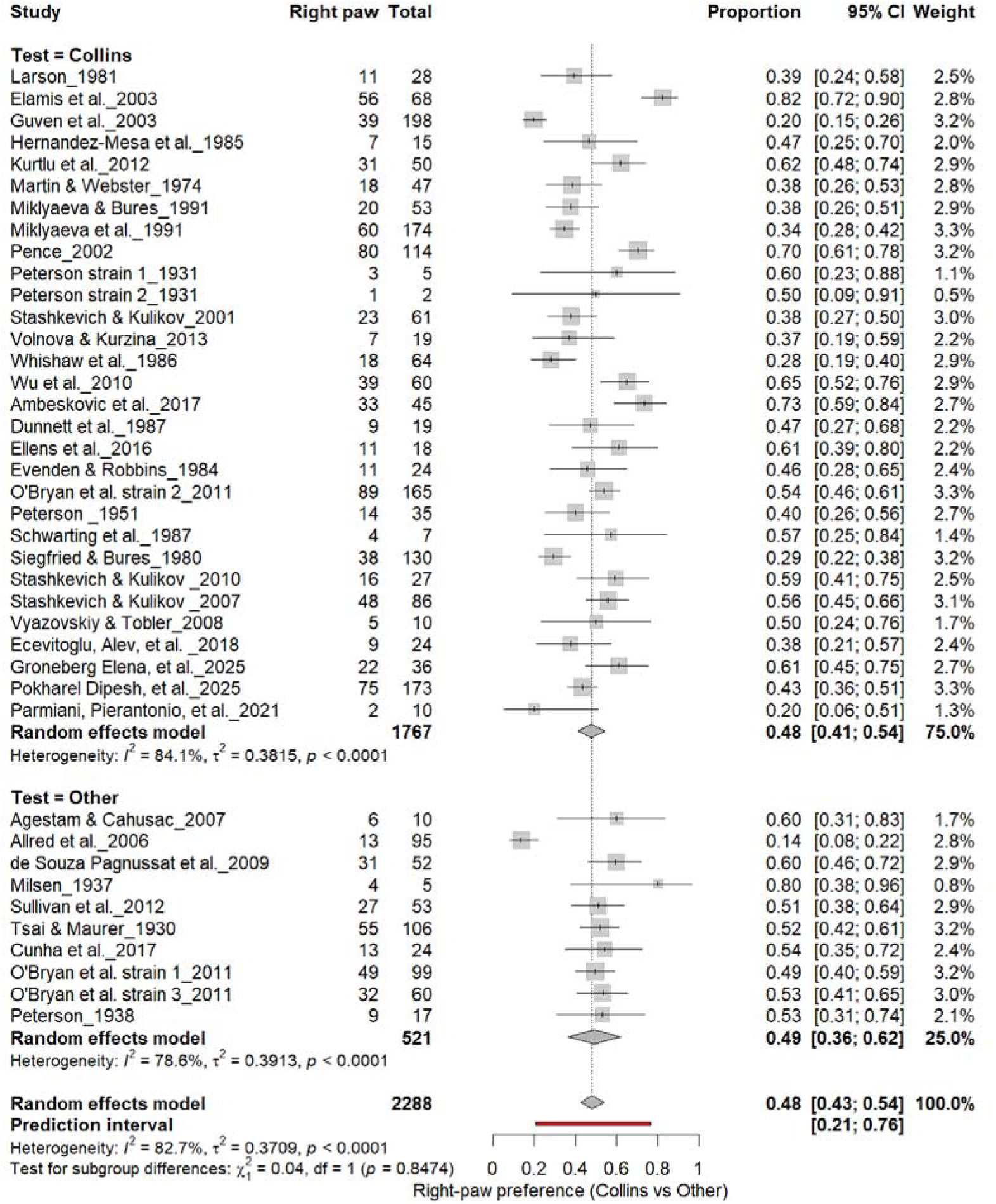
Forest plot (n = 2,288 rats) showing the prevalence of right-paw preference. Each square represents a study-specific estimate (size proportional to study weight), with horizontal lines denoting 95% confidence intervals. The pooled estimate from a random-effects model indicates that 48% of rats were right-pawed (95% CI: 0.43–0.54, p < 0.0001). Substantial between-study heterogeneity was observed (I² = 82.7%), with no significant difference between the Collins test and other behavioral paradigms (χ² = 0.04, df = 1, p = 0.85).

**Figure 8.**
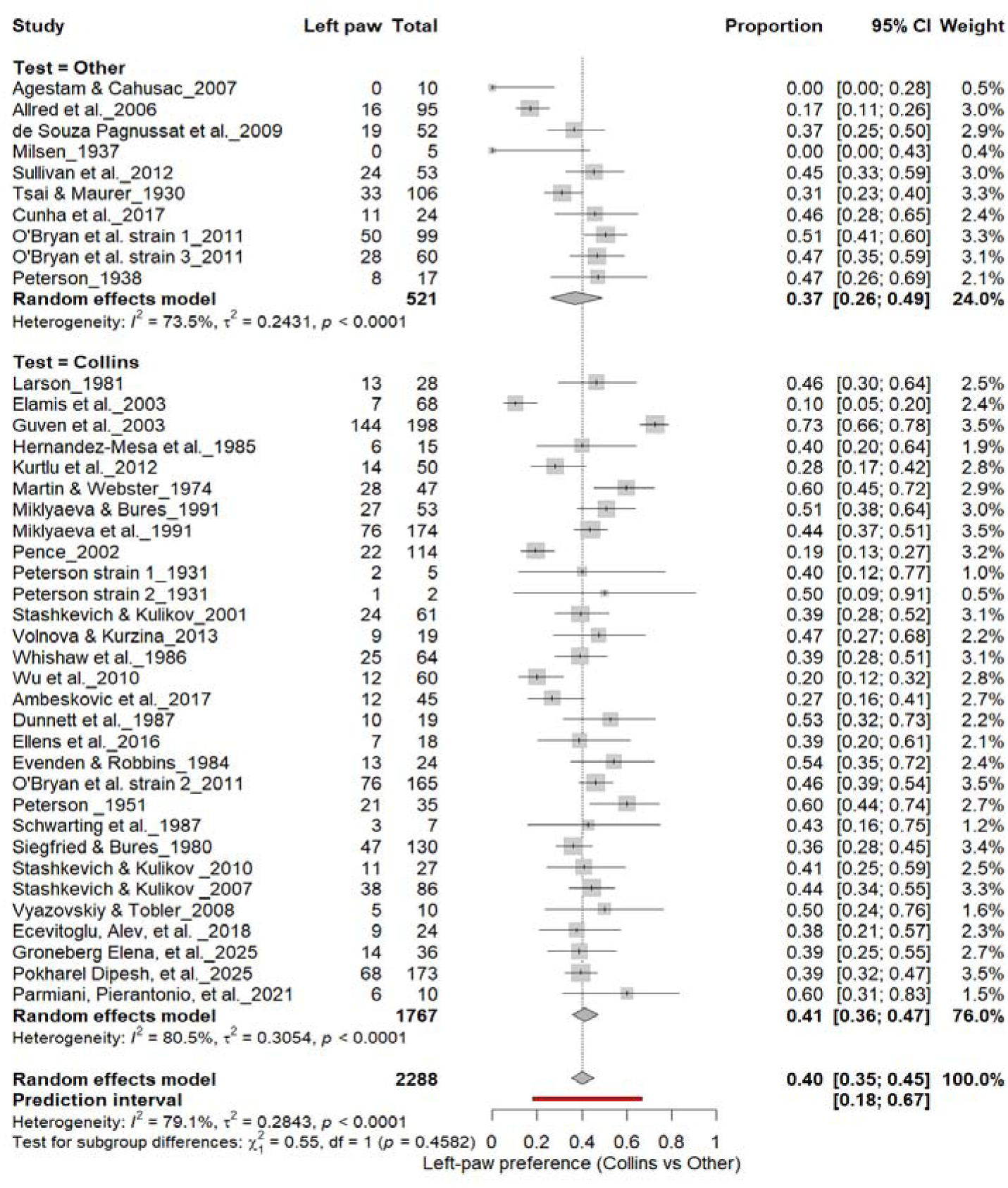
Forest plot (n = 2,288 rats) showing the prevalence of left-paw preference. Each square represents a study-specific estimate (size proportional to study weight), and horizontal lines denote 95% confidence intervals. The pooled estimate from a random-effects model indicates that 40% of rats were left-pawed (95% CI: 0.35–0.45, p < 0.0001). Substantial between-study heterogeneity was observed (I² = 79.1%), with no significant difference between the Collins test and other behavioral paradigms (χ² = 0.28, df = 1, p = 0.60).

**Figure 9.**
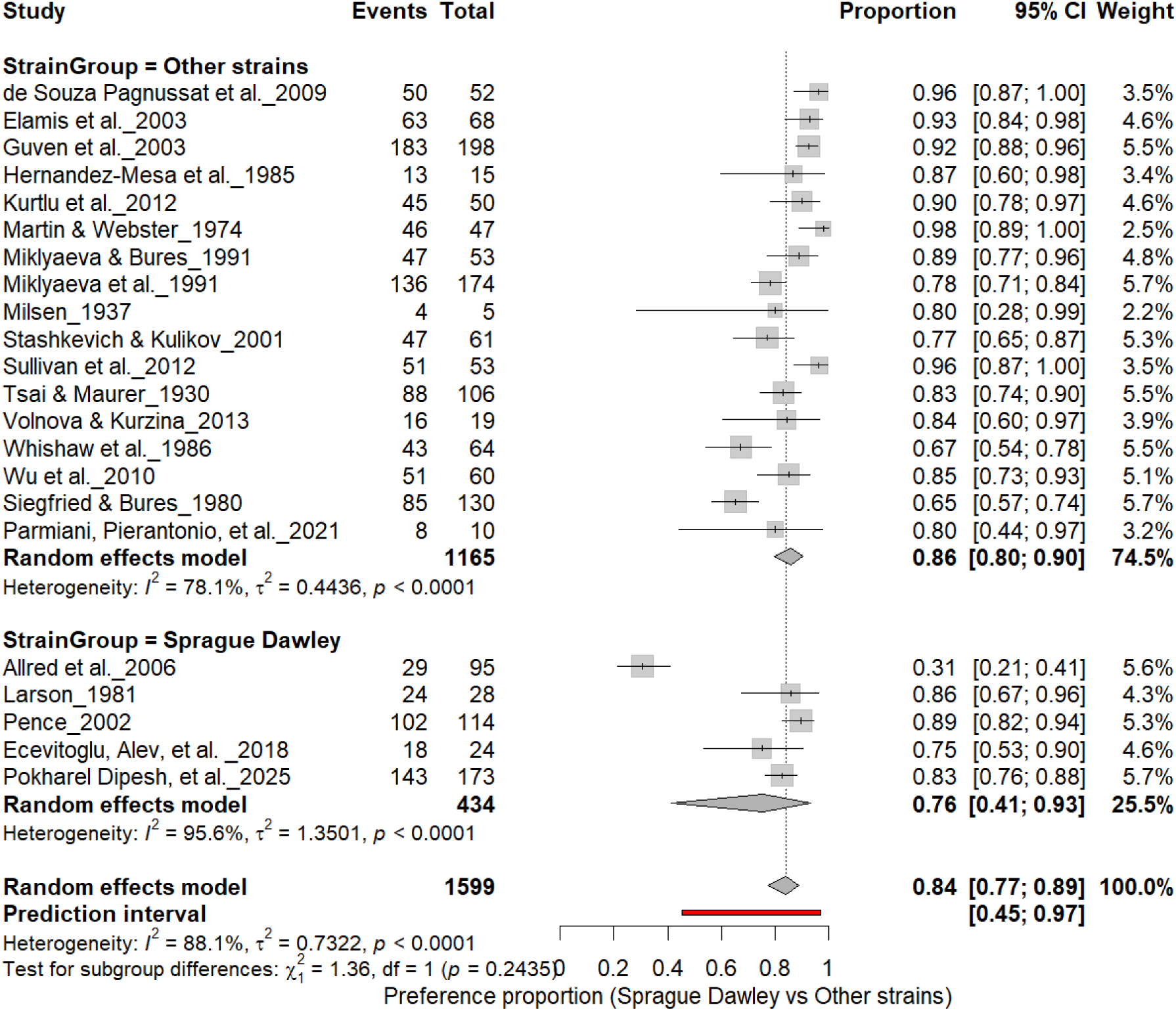
Forest plot (n = 1,599 rats) comparing paw preference prevalence between Sprague Dawley and other rat strains. Each square represents a study-specific estimate (size proportional to study weight), with horizontal lines indicating 95% confidence intervals. The pooled estimate from a random-effects model shows that 84% of rats exhibited consistent paw preference (95% CI: 0.77–0.89, p < 0.0001). Subgroup analysis revealed prevalence rates of 76% in Sprague Dawley and 86% in other strains, with no significant subgroup difference (χ² = 1.36, df = 1, p = 0.24).

**Figure 10.**
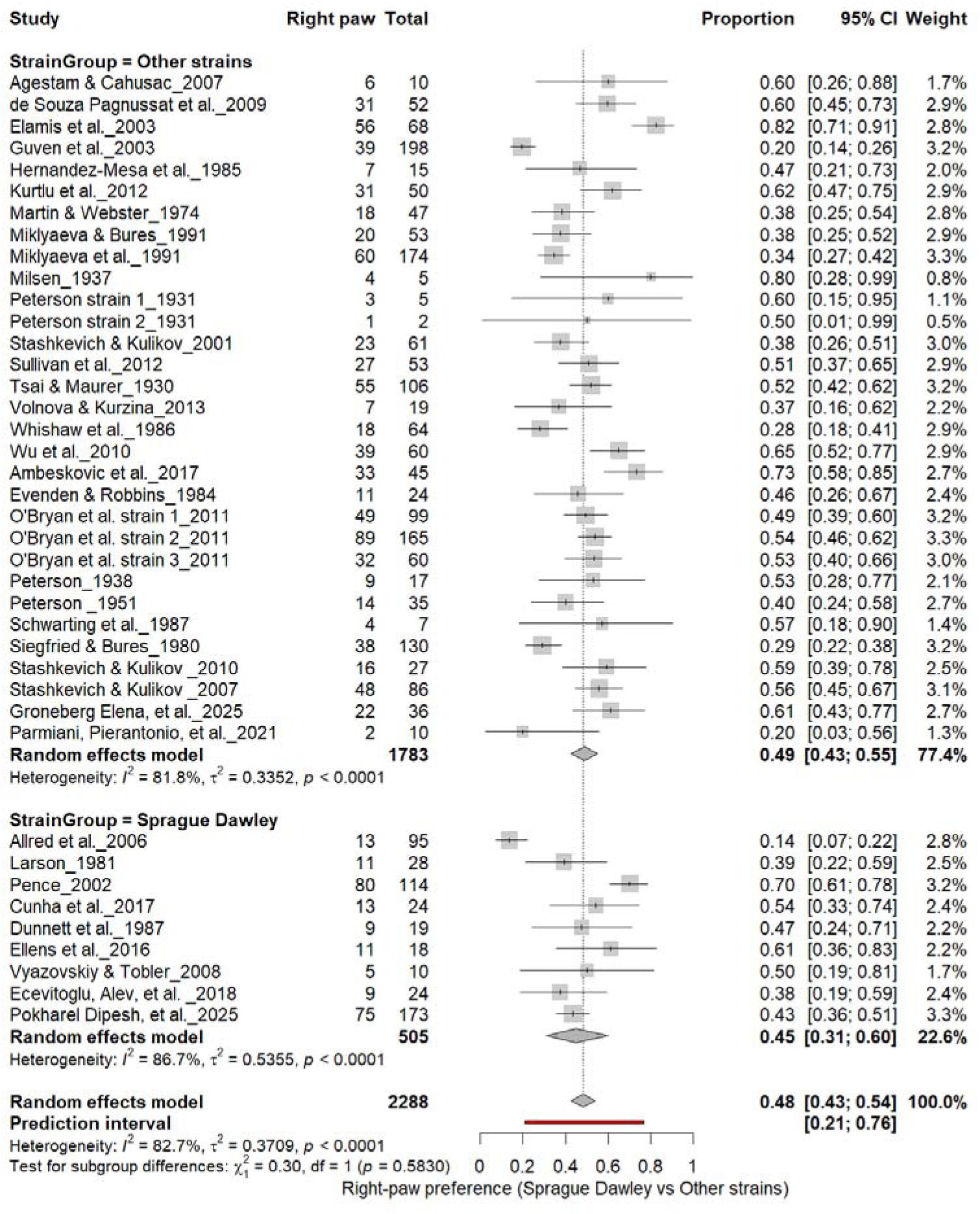
Forest plot (n = 2,288 rats) showing the prevalence of right-paw preference across strains. Each square represents a study-specific estimate (size proportional to study weight), and horizontal lines indicate 95% confidence intervals. The pooled estimate from a random-effects model indicates that 48% of rats were right-pawed (95% CI: 0.43–0.54, p < 0.0001). Subgroup analysis revealed no significant difference between Sprague Dawley and other strains (χ² = 0.30,

**Figure 11.**
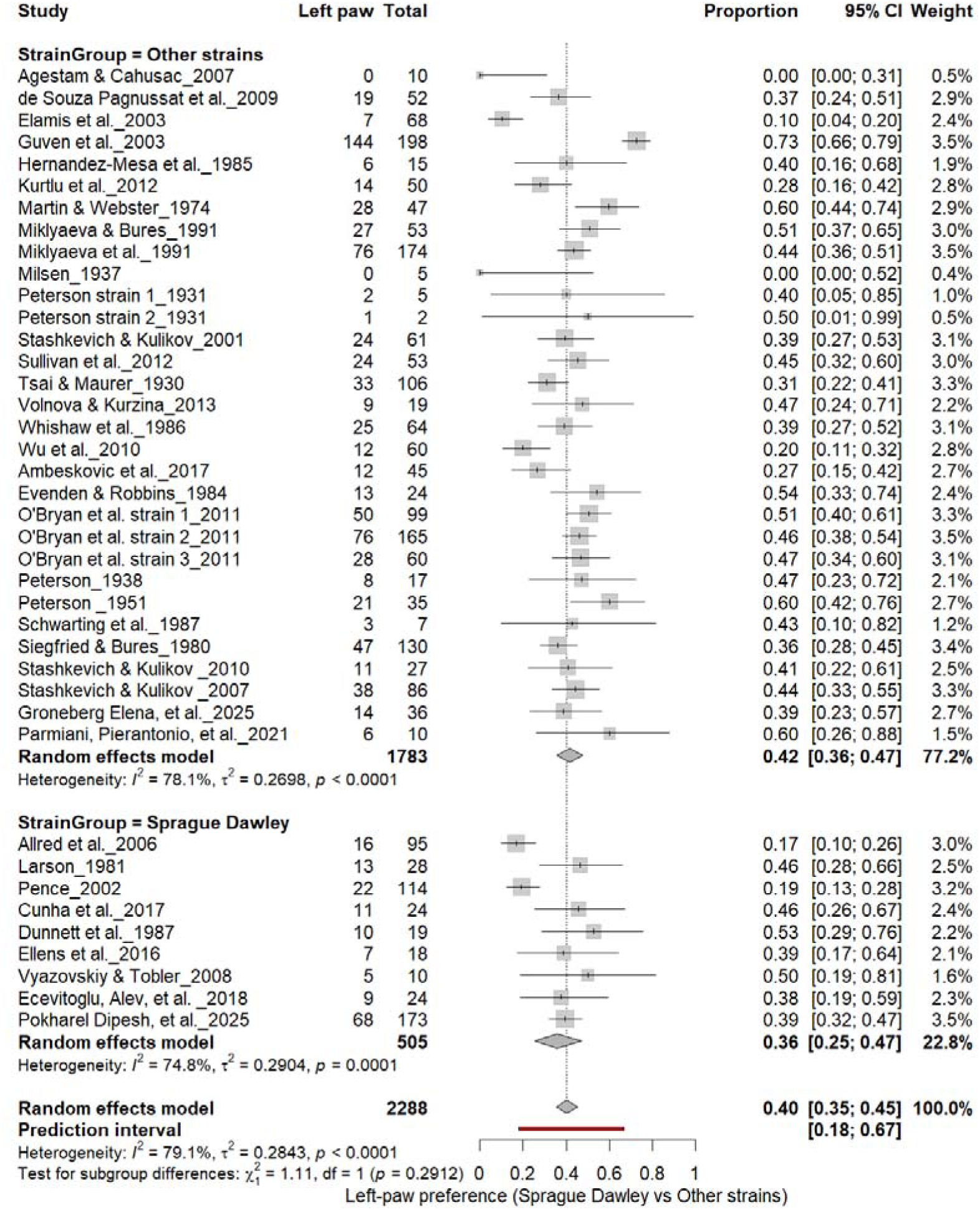
Forest plot (n = 2,288 rats) showing the prevalence of left-paw preference across strains. Each square represents a study-specific estimate (size proportional to study weight), and horizontal lines denote 95% confidence intervals. The pooled estimate from a random-effects model indicates that 40% of rats were left-pawed (95% CI: 0.35–0.45, p < 0.0001). Subgroup analysis revealed no significant difference between Sprague Dawley and other strains (χ² = 1.11,

**Figure 12.**
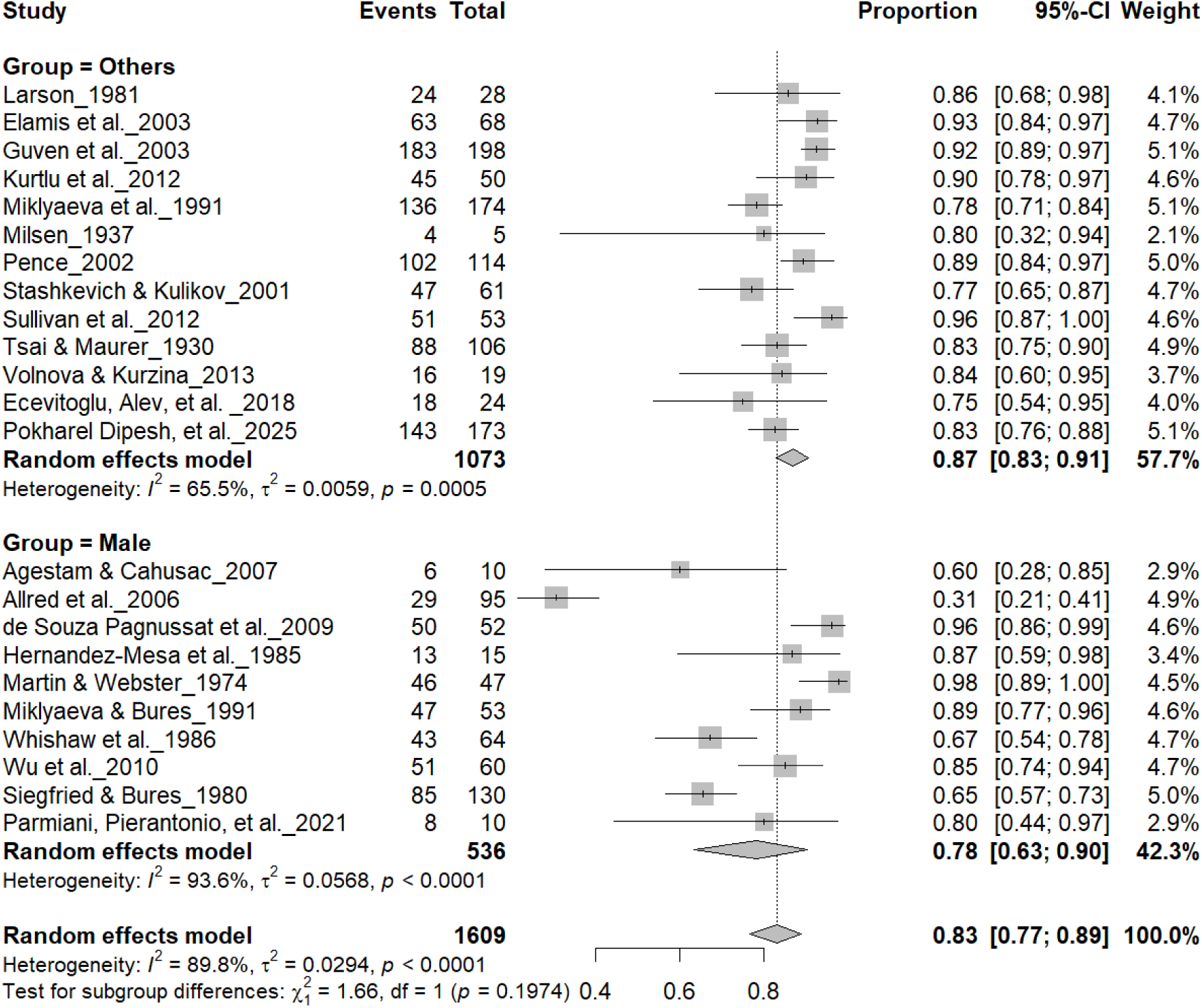
Forest plot (n = 1,609 rats) showing consistent paw preference by sex. Each square represents a study-specific estimate (size proportional to study weight), with horizontal lines denoting 95% confidence intervals. The pooled estimate from a random-effects model indicates that 83% of rats exhibited consistent paw preference (95% CI: 0.77–0.89, p < 0.0001). Subgroup analysis showed similar prevalence between males (83%) and other groups, with no significant difference (χ² = 1.17, df = 1, p = 0.28).

**Figure 13.**
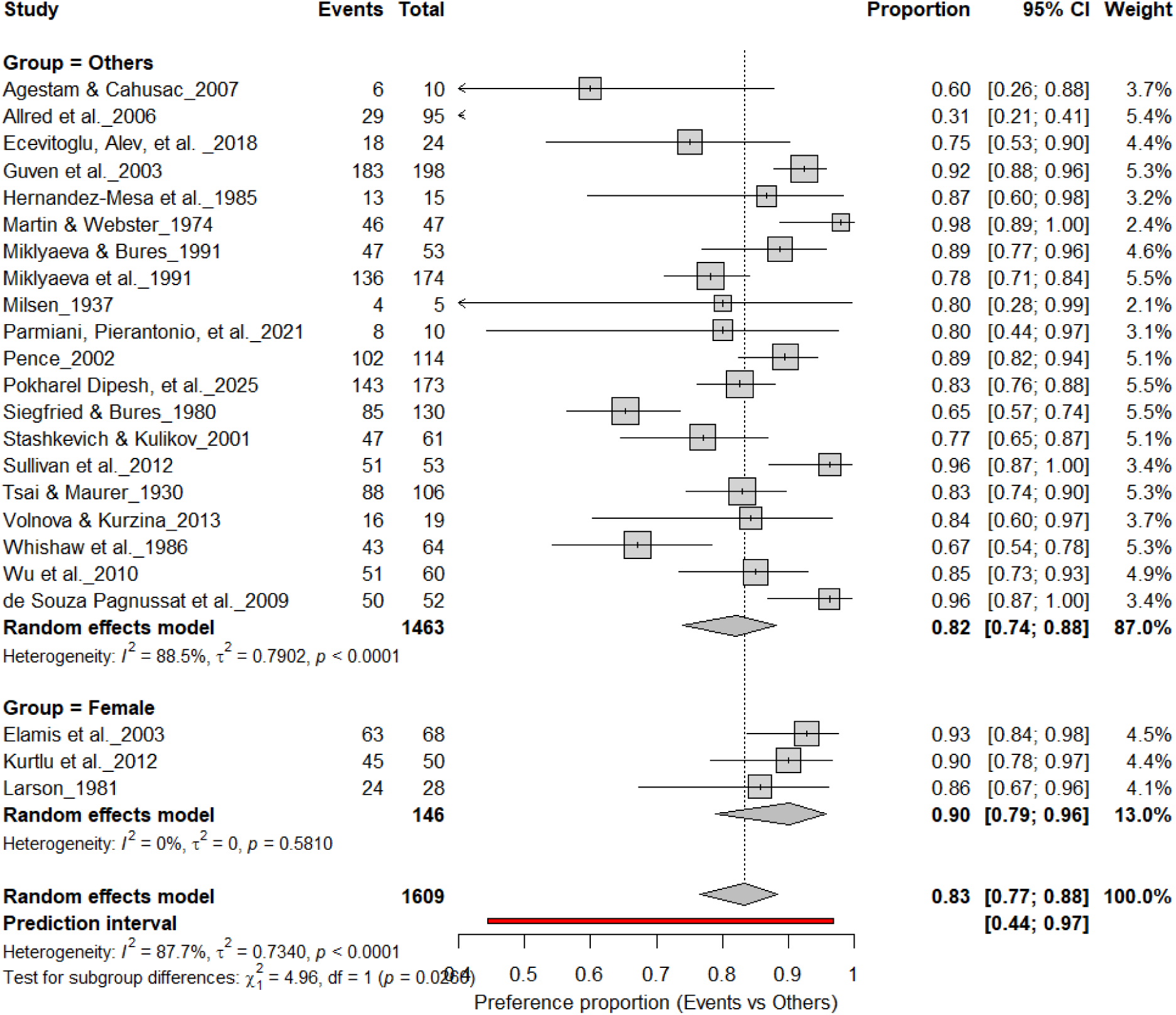
Forest plot (n = 1,690 rats) comparing paw preference prevalence between female rats and other groups. Each square represents a study-specific estimate (size proportional to study weight), with horizontal lines indicating 95% confidence intervals. The pooled estimate from a random-effects model indicates that 83% of rats exhibited consistent paw preference (95% CI: 0.77–0.89, p < 0.0001). Subgroup analysis showed similar prevalence between females (83%) and other groups (82%), with no significant difference (χ² = 1.48, df = 1, p = 0.22).

**Figure 14.**
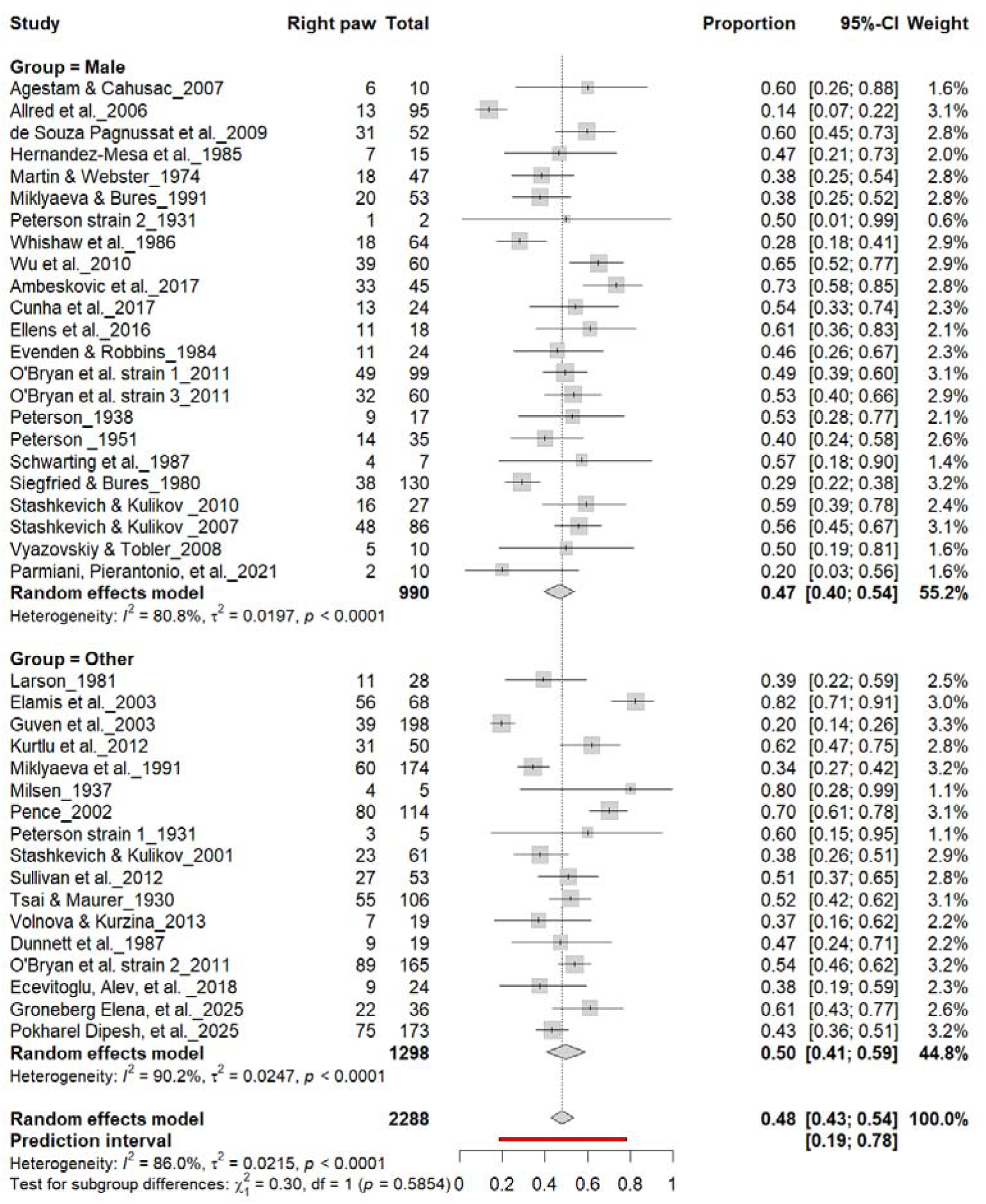
Forest plot (n = 2,288 rats) showing the prevalence of right-paw preference by sex. Each square represents a study-specific estimate (size proportional to study weight), with horizontal lines indicating 95% confidence intervals. The pooled estimate from a random-effects model indicates that 48% of rats were right-pawed (95% CI: 0.43–0.54, p < 0.0001). Subgroup analysis revealed similar prevalence between males (47%) and other groups (45%), with no significant difference (χ² = 0.03, df = 1, p = 0.56).

**Figure 15.**
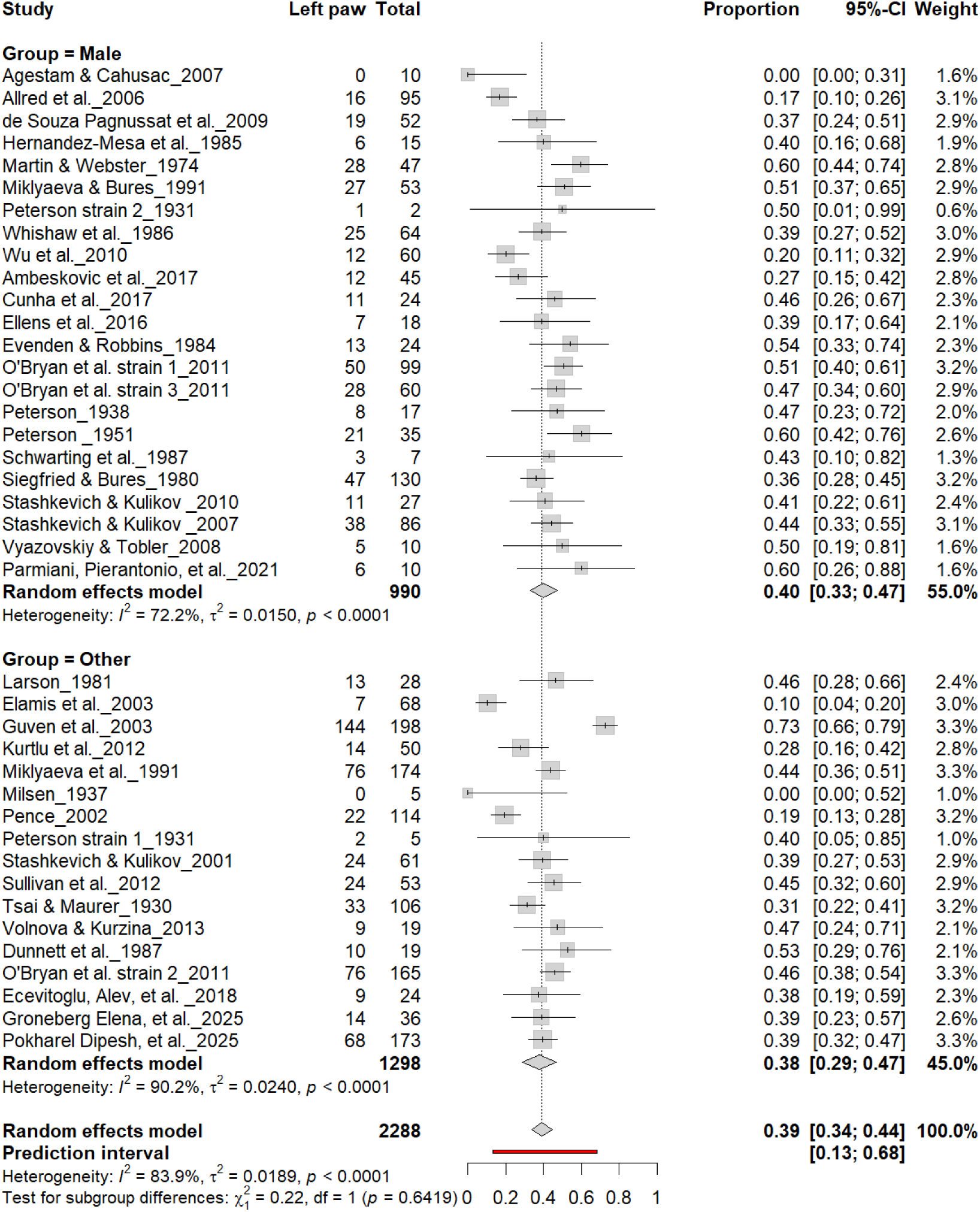
Forest plot (n = 2,288 rats) showing the prevalence of left-paw preference by sex. Each square represents a study-specific estimate (size proportional to study weight), and horizontal lines denote 95% confidence intervals. The pooled estimate from a random-effects model indicates that 39% of rats were left-pawed (95% CI: 0.34–0.44, p < 0.0001). Subgroup analysis revealed similar prevalence between males (40%) and other groups (38%), with no significant difference (χ² = 0.22, df = 1, p = 0.64).

**Figure 16.**
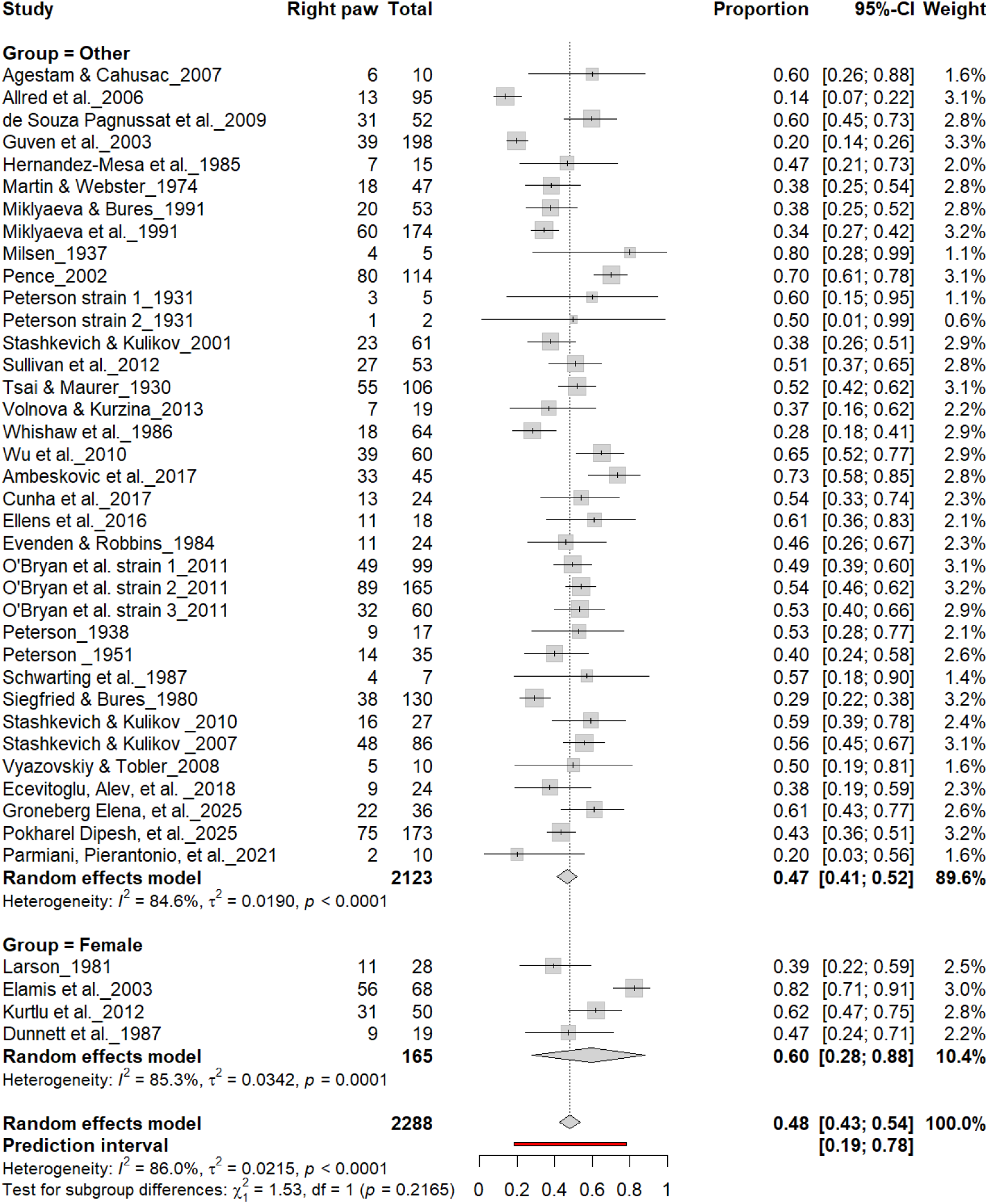
Forest plot (n = 2,288 rats) showing the prevalence of right-paw preference by sex. Each square represents a study-specific estimate (size proportional to study weight), and horizontal lines indicate 95% confidence intervals. The pooled estimate from a random-effects model indicates that 48% of rats were right-pawed (95% CI: 0.43–0.54, p < 0.0001). Subgroup analysis revealed no significant difference between female rats (60%) and other groups (47%)

**Figure 17.**
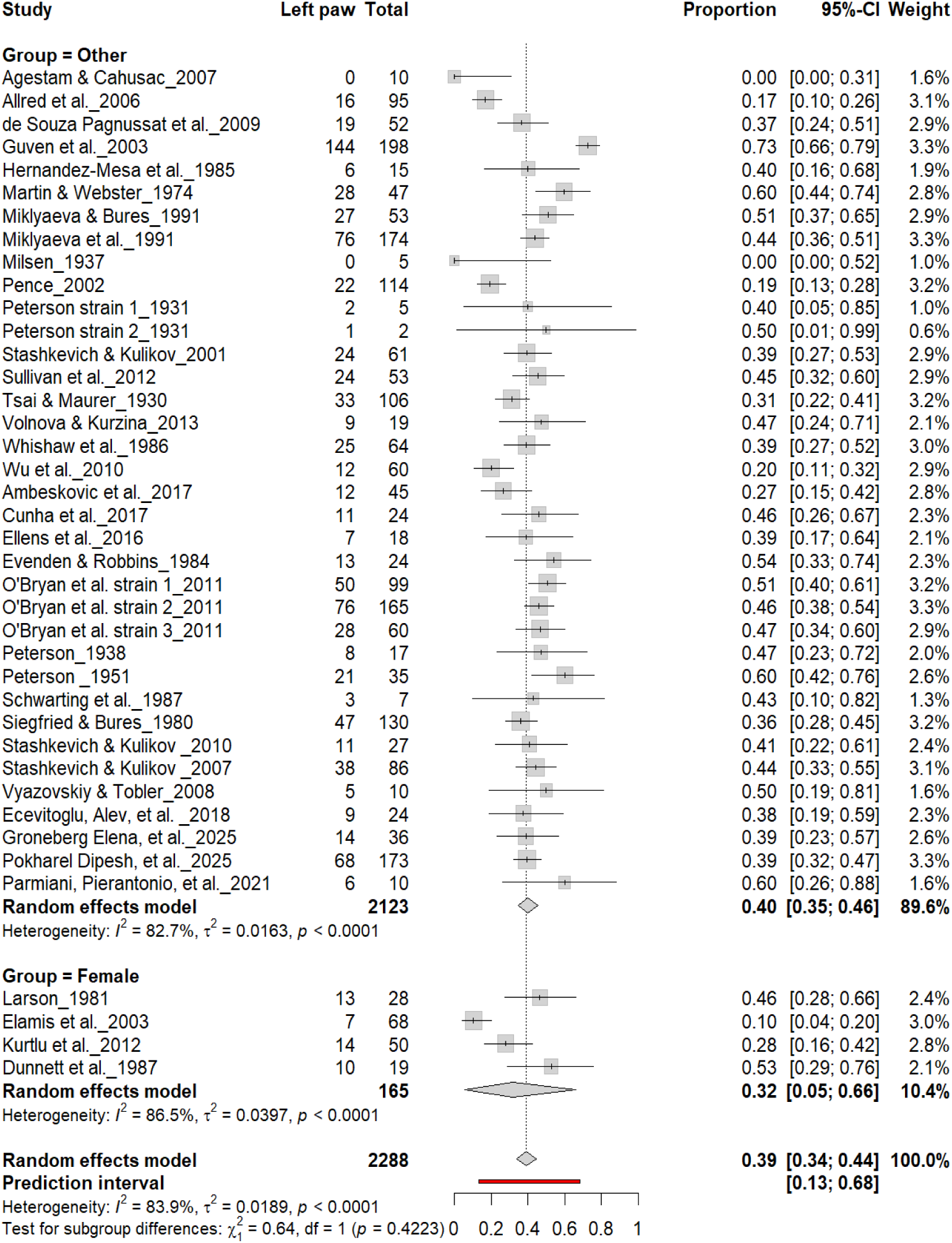
Forest plot (n = 2,288 rats) showing the prevalence of left-paw preference by sex. Each square represents a study-specific estimate (size proportional to study weight), and horizontal lines denote 95% confidence intervals. The pooled estimate from a random-effects model indicates that 39% of rats were left-pawed (95% CI: 0.34–0.44, p < 0.0001). Subgroup analysis revealed no significant difference between female rats (32%) and other groups (40%)

**Figure 18.**
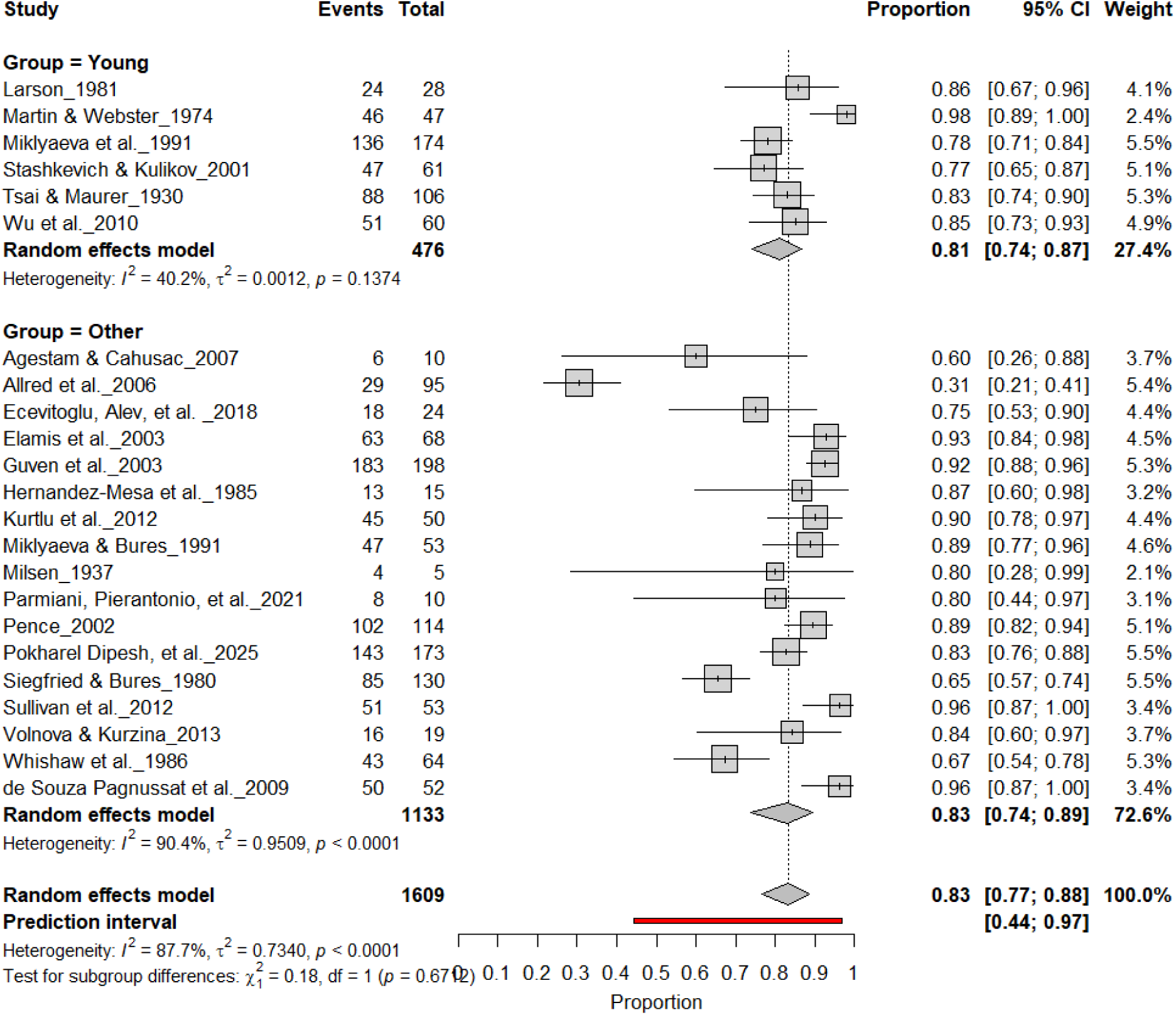
Forest plot (n = 1,609 rats) showing consistent paw preference prevalence across age groups. Each square represents a study-specific estimate (size proportional to study weight), and horizontal lines denote 95% confidence intervals. The pooled estimate from a random-effects model indicates that 83% of rats exhibited consistent paw preference (95% CI: 0.77–0.88, p < 0.0001). Subgroup analysis revealed similar prevalence in both young and adult rats, with no significant age-related difference (χ² = 0.18, df = 1, p = 0.67).

**Figure 19.**
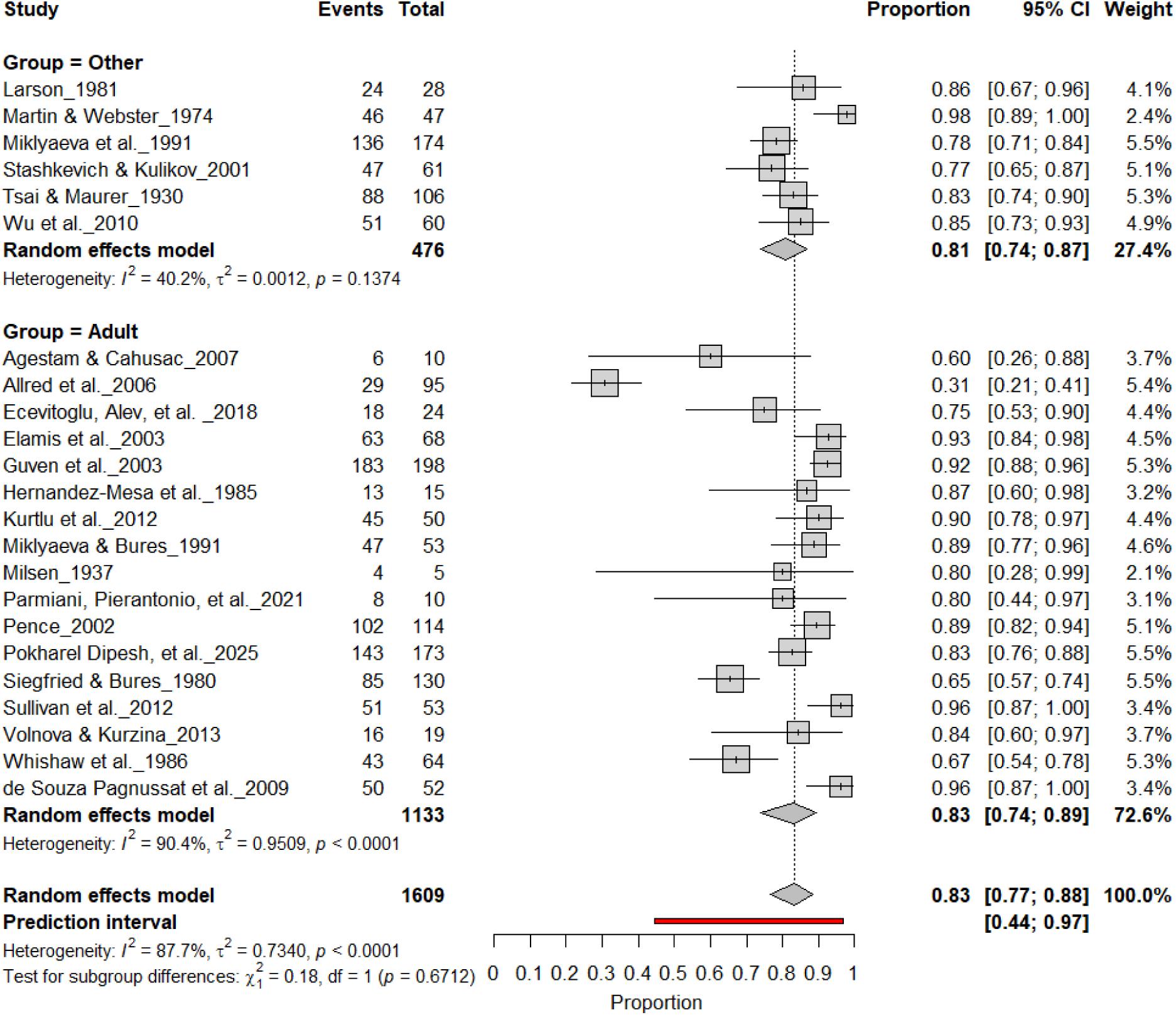
Forest plot of (n = 1,609 rats) showing consistent paw preference prevalence across age groups. Each square represents a study-specific estimate (size proportional to study weight), and horizontal lines denote 95% confidence intervals. The pooled estimate from a random-effects model indicates that 83% of rats exhibited consistent paw preference (95% CI: 0.77–0.88, p < 0.0001). Subgroup analysis revealed comparable prevalence between adult and younger rats, with no significant subgroup difference (χ² = 0.18, df = 1, p = 0.67).

**Figure 20.**
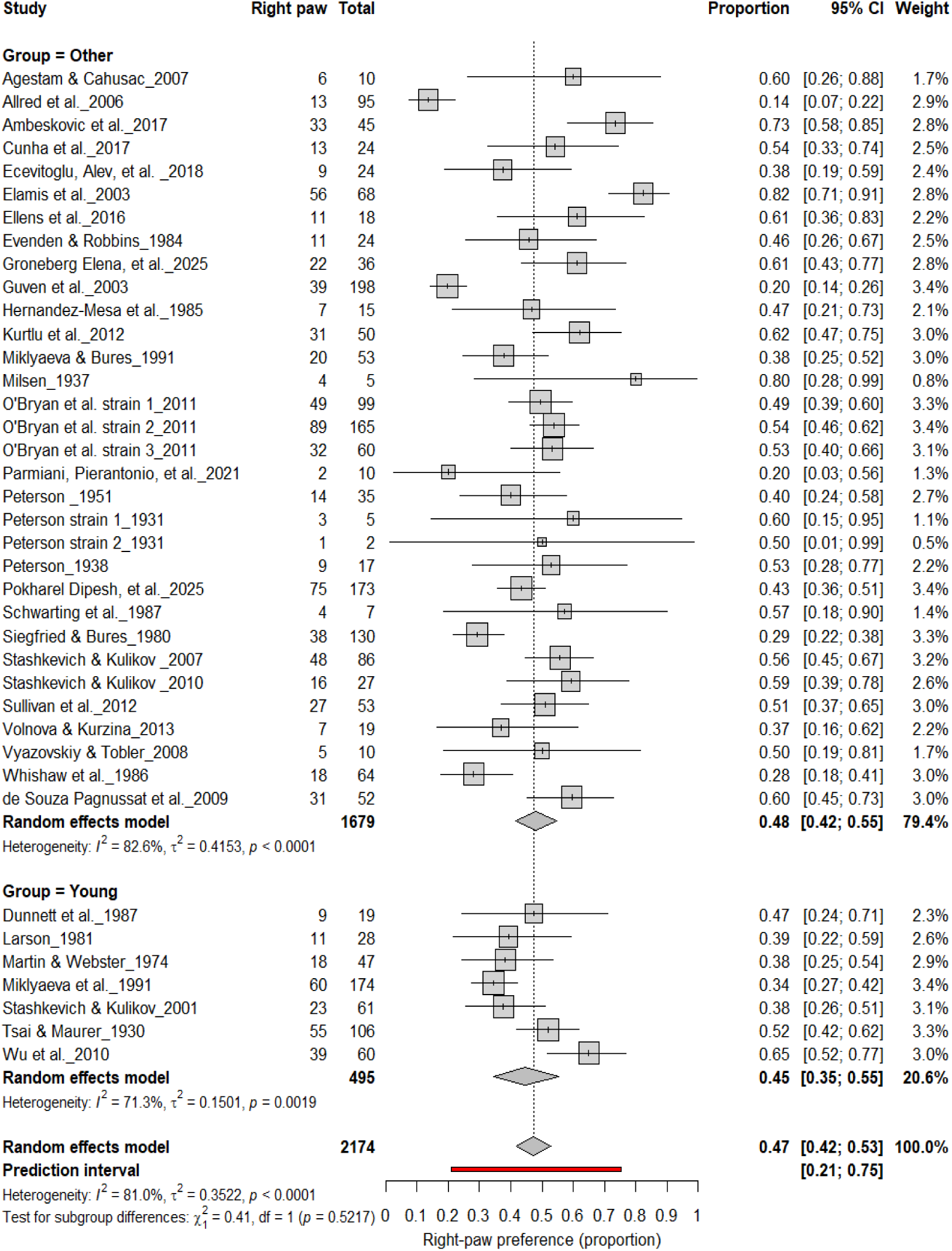
Forest plot (n = 2,174 rats) showing the prevalence of right-paw preference by age group. Each square represents a study-specific estimate (size proportional to study weight), and horizontal lines indicate 95% confidence intervals. The pooled estimate from a random-effects model indicates that 47% of rats were right-pawed (95% CI: 0.42–0.53, p < 0.0001). Subgroup analysis revealed comparable prevalence between young (45%) and older rats (48%), with no significant difference (χ² = 0.41, df = 1, p = 0.52).

**Figure 21.**
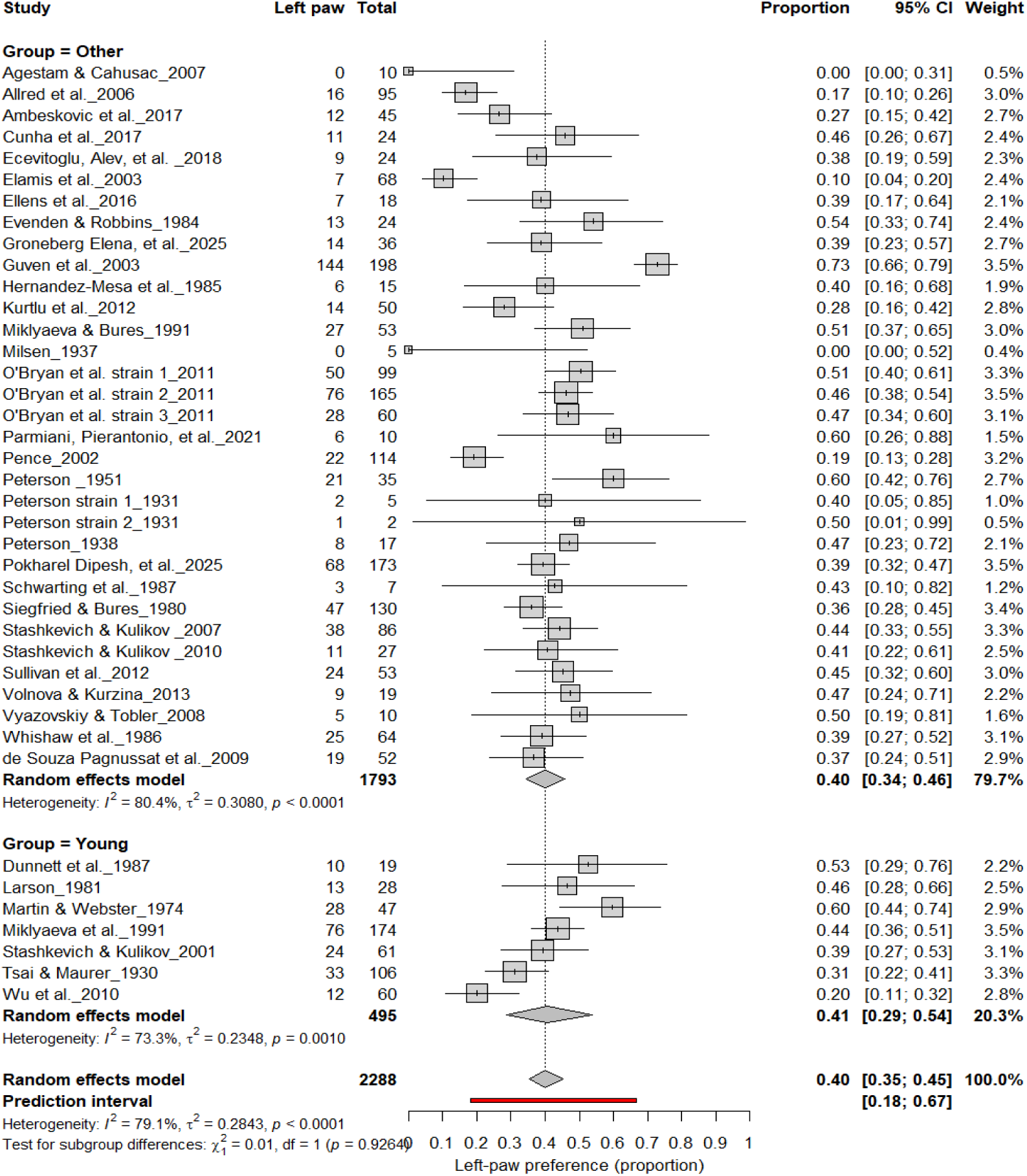
Forest plot (n = 2,288 rats) showing the prevalence of left-paw preference by age group. Each square represents a study-specific estimate (size proportional to study weight), with horizontal lines indicating 95% confidence intervals. The pooled estimate from a random-effects model indicates that 40% of rats were left-pawed (95% CI: 0.35–0.45, p < 0.0001). Subgroup analysis revealed comparable prevalence between young and older rats, with no significant difference (χ² = 2.00, df = 1, p = 0.16).

**Figure 22.**
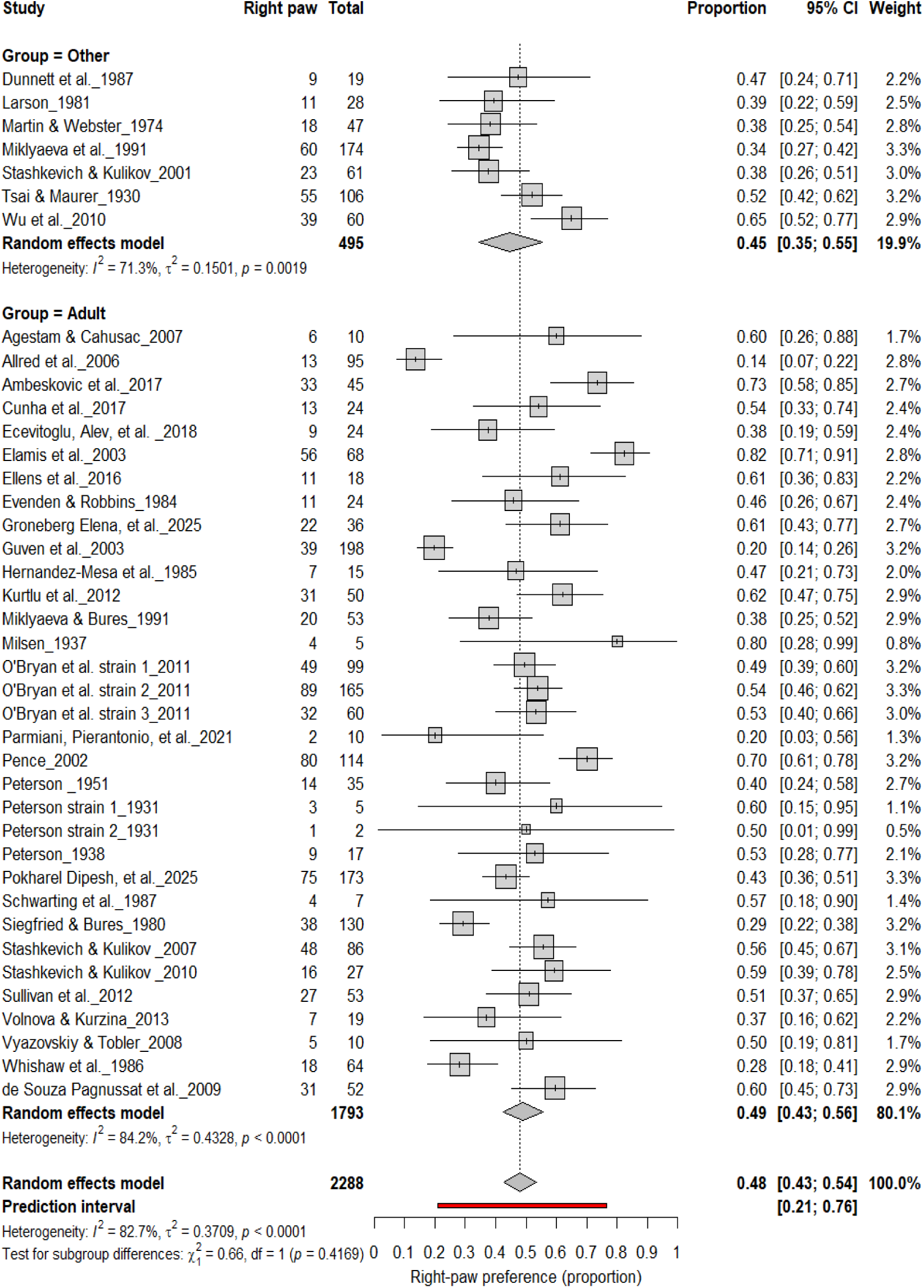
Forest plot of (n = 2,288 rats) showing the prevalence of right-paw preference by age group. Each square represents a study-specific estimate (size proportional to study weight), and horizontal lines indicate 95% confidence intervals. The pooled estimate from a random-effects model indicates that 48% of rats were right-pawed (95% CI: 0.43–0.54, p < 0.0001). Subgroup analysis revealed no significant difference between adult and other rats (χ² = 0.01, df = 1, p = 0.91).

**Figure 23.**
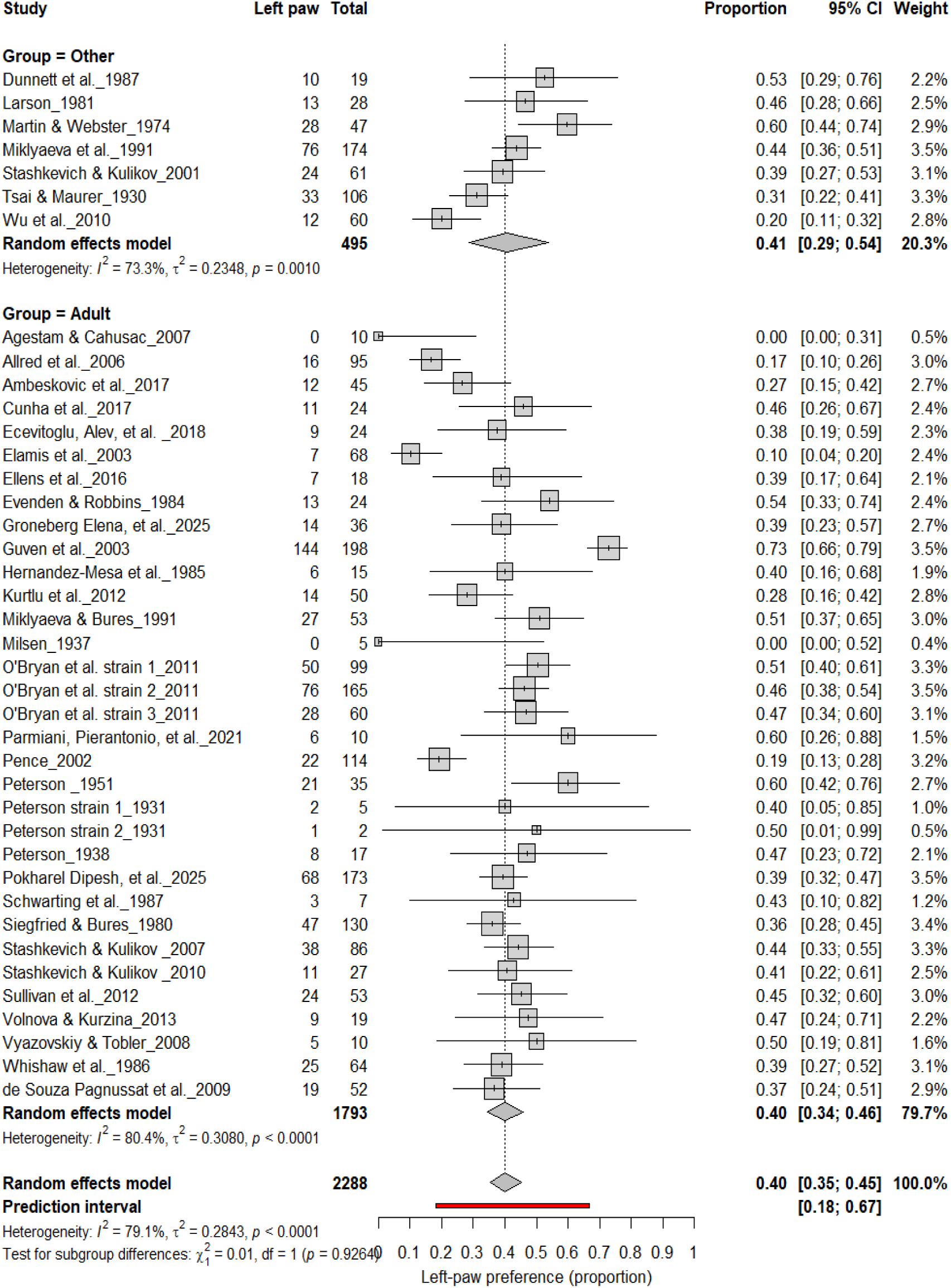
Forest plot (n = 2,288 rats) showing the prevalence of left-paw preference by age group. Each square represents a study-specific estimate (size proportional to study weight), with horizontal lines indicating 95% confidence intervals. The pooled estimate from a random-effects model indicates that 40% of rats were left-pawed (95% CI: 0.35–0.45, p < 0.0001). Subgroup analysis revealed comparable prevalence between adult and other rats, with no significant difference (χ² = 2.00, df = 1, p = 0.16).

### 2.10 Statistical synthesis methods

Random-effects models were applied to logit-transformed event rates to account for between-study variability, yielding prevalence estimates and 95% confidence intervals for (i) lateralization (left + right vs. ambidextrous) and (ii) directional bias (right vs. non-right; left vs. non-left). Heterogeneity (Q, I², τ²) and 95% prediction intervals were calculated, and subgroup analyses examined strain, sex, age, and test paradigm effects.

### 2.11 Sensitivity Analyses

Sensitivity analyses were performed to evaluate the robustness of results. Studies assessed as having high risk of bias on the SYRCLE tool were sequentially excluded to determine their impact on pooled estimates.

### 2.12 Heterogeneity

Heterogeneity was assessed using Cochran’s Q statistic, the I² index, and the between-study variance estimate (τ²). I² values of 25%, 50%, and 75% were interpreted as low, moderate, and high heterogeneity, respectively. Event rates were transformed using logit proportions, and all pooled estimates were reported with 95% confidence intervals (CIs).^57^ Comparisons against chance-level preference (0.5) were tested using Z-tests.

### 2.13 Software

All statistical analyses were performed using R (version 4.2.0; R Foundation for Statistical Computing, Vienna, Austria) via RStudio (version 2025.09.0; Posit Software, PBC). Forest plots were generated using the *meta* package. The *metaprop* function was used to compute pooled prevalence estimates, while the *metabin* function was applied for subgroup comparisons.^58^

## 3. Results

### 3.1 Study characteristics

This meta-analysis included 40 studies published between 1930 and 2025 that evaluated paw preference in rats, encompassing a total of 1,609 animals (Table 1). Rat strains were diverse, with Sprague Dawley rats being the most frequently studied (17/40 studies), followed by Wistar rats (13/40), while other strains such as Long-Evans, Lister Hooded, Hooded Druckray, and Albino were less commonly used. Sex distribution varied considerably across studies. Out of 85 experimental groups, 30 included only males, 18 included only females, 28 included both sexes, and 9 did not specify sex. Similarly, age was not uniformly reported, with 47 groups studying adult rats, 12 young rats, 3 mixed-age cohorts, and 23 groups in which age was not clearly specified. A range of behavioral assays were employed to assess paw preference. The Collins paw preference test and its modifications were the most frequently used paradigm (17/40 studies), while skilled reaching tasks including the single pellet reaching test, staircase test, and vermicelli handling test were reported in 15/40 studies. Other less common approaches included food-reaching and board tasks, circular cage tests, and additional methods such as tube food-reaching, and lever-pressing tasks.

### 3.2 Study Risk of Bias Analysis

The risk of bias of studies was assessed with the SYRCLE tool, adapted for this systematic review. 10 questions were answered for each study with an answer of “yes” indicating lower risk of bias and an answer of “unclear” or “no” indicating a higher risk of bias (Table 2). The results indicate variable bias amongst the studies. It was noted that many studies are lacking in complete transparency of these details, leading to many answers of “unclear”. Questions that received more “unclear” and “no” answers, indicating higher risk of bias were 1,3,5,6, and 7. Questions that received more “yes” answers, indicating lower risk of bias were 2,4,8,9, and 10. Overall, transparency and methodological rigor were inconsistent across the dataset, which may contribute to observed heterogeneity in pooled outcomes (Figure 2; Table 2).

### 3.3 Individual-level Paw Preference Asymmetries

Figure 3 summarizes 24 studies (n = 1,609 rats) assessing the prevalence of consistent paw preference. Each square represents the proportion of rats exhibiting stable lateralization, with square size proportional to study weight and horizontal lines denoting 95% confidence intervals (CI). The pooled estimate from a random-effects model indicated that 83% of rats exhibited consistent paw preference (95% CI: 0.77–0.88, *p* < 0.0001), with a 95% prediction interval of 0.44–0.97, reflecting considerable inter-study variability. Despite substantial heterogeneity (I² = 87.7%), these findings demonstrate robust and reproducible individual-level lateralization across rat strains and behavioral paradigms.

### 3.4 Population-Level Paw Preference Asymmetries

#### 3.4A. Right-Pawedness vs. Non-Right-Pawedness

Figure 4 summarizes 40 studies (n = 2,288 rats) evaluating the proportion of right-pawed rats. The pooled estimate from a random-effects model indicated that 48% of rats were right-pawed (95% CI: 43–54%, *p* < 0.0001), with substantial heterogeneity (I² = 82.7%) and a prediction interval of 21–76%, suggesting considerable between-study variability. These findings indicate no consistent population-level rightward bias, reinforcing that lateralization in rats primarily occurs at the individual rather than population level.

#### 3.4 B. Left-Pawedness vs. Non-Left-Pawedness

Figure 5 summarizes 40 studies (n = 2,288 rats) evaluating the proportion of left-pawed rats. The pooled estimate from a random-effects model indicated that 40% of rats were left-pawed (95% CI: 35–45%), with substantial heterogeneity (I² = 79.1%) and a prediction interval of 18–67%, reflecting notable between-study variability. As with right-pawedness, these results demonstrate no consistent population-level leftward bias, supporting that forelimb lateralization in rats emerges at the individual rather than population level.

### 3.5 Test Paradigm Effects

#### 3.5 A. Individual-Level Paw Preference: Collins Test vs. Other Paradigms

Figure 6 compares Collins-based tests (17 cohorts, *n* = 1,288) with other paradigms (6 cohorts, *n* = 321) for assessing individual-level paw preference. Both approaches demonstrated strong and comparable lateralization: Collins test: 83% (*95% CI*: 79–88%, *I²* = 76.1%) and other paradigms: 81% (*95% CI*: 43–96%, *I²* = 93.9%). The overall pooled prevalence across all cohorts was 83% (*95% CI*: 77–88%), with a prediction interval of 44–97%, reflecting substantial inter-study variability. Subgroup comparison was not significant (χ² = 0.11, df = 1, *p* = 0.74), confirming that the strength of individual-level asymmetry does not differ between test paradigms.

### 3.6 Population-Level Asymmetries by Test Paradigm

#### 3.6 A. Right-Pawedness: Collins vs. Other Tests

Figure 7 summarizes (n = 2,288 rats). The Collins test yielded a pooled prevalence of 48% (95% CI: 41–54%), while other paradigms produced 48% (95% CI: 31–66%). Heterogeneity was high in both subgroups (I² = 75.0%–78.0%) and overall (I² = 82.7%). Subgroup differences were not significant (χ² = 0.04, df = 1, p = 0.85). The prediction interval (21–77%) indicates broad inter-study variability, confirming that task type does not affect right-paw prevalence.

#### 3.6 B. Left-Pawedness: Collins vs. Other Tests

Figure 8 shows comparable findings. The Collins test yielded 40% (95% CI: 35–45%), while other paradigms produced 47% (95% CI: 29–64%), with substantial heterogeneity (I² = 79.1%). The overall pooled prevalence was 40% (95% CI: 35–45%), and subgroup comparison remained non-significant (χ² = 0.28, df = 1, p = 0.60). Thus, the testing paradigm did not influence population-level asymmetry, confirming that leftward bias was not systematically affected by task type.

### 3.7 Strain Effects

#### 3.7 A. Individual-Level Paw Preference: Sprague Dawley vs. Other Strains

Figure 9 summarizes (n = 1,599 rats) comparing paw preference prevalence across strains. Non–Sprague Dawley cohorts (17 cohorts, n = 1,165) showed a pooled prevalence of 86% (95% CI: 79–90%), whereas Sprague Dawley cohorts (5 cohorts, n = 434) showed 76% (95% CI: 54–89%). Heterogeneity was substantial across both subgroups (I² = 78.1%–95.6%), and the subgroup difference was not significant (χ² = 1.36, df = 1, p = 0.24). Overall, strain background modestly influenced strength but not the direction of asymmetry, suggesting robust individual-level lateralization across genetic backgrounds.

### 3.8 Population-Level Paw Preference: Sprague Dawley vs. Other Strains

#### 3.8 A. Right-Pawedness: Sprague Dawley vs. Other Strains

Figure 10 summarizes (n = 2,288 rats) comparing right-paw prevalence between Sprague Dawley and other rat strains. Non–Sprague Dawley cohorts showed a pooled prevalence of 49% (95% CI: 43–54%), while Sprague Dawley rats showed 45% (95% CI: 33–58%), with high heterogeneity across subgroups (I² = 81.8%–85.7%). The overall pooled prevalence was 48% (95% CI: 43–54%), and subgroup differences were not significant (χ² = 0.30, df = 1, p = 0.58). These findings indicate no strain-dependent rightward bias in paw preference.

#### 3.8 B. Left-Pawedness: Sprague Dawley vs. Other Strains

Figure 11 compares left-paw prevalence across rat strains. Non–Sprague Dawley cohorts showed 40% (95% CI: 35–44%), whereas Sprague Dawley cohorts showed 35% (95% CI: 26–45%), both exhibiting high heterogeneity (I² = 74.6%–77.1%). The overall pooled prevalence was 39% (95% CI: 34–44%), and subgroup comparison was not significant (χ² = 1.11, df = 1, p = 0.29). These findings indicate that strain background did not influence the direction of population-level asymmetry, reaffirming the absence of consistent leftward bias across strains.

### 3.9 Sex Effects

#### 3.9 A. Individual-Level Asymmetry (Male and Female)

Figures 12 and 13 summarize (n = 1,497 rats) comparing consistent paw preference between male and female rats. Males exhibited a pooled prevalence of 83% (95% CI: 77–89%), while females showed 92% (95% CI: 85–95%). Heterogeneity was lower in females (I² = 9.5%) than in males (I² = 63.8%). Although females appeared slightly more lateralized, subgroup analysis revealed no significant difference between sexes (χ² = 1.48, df = 1, p = 0.22). Both sexes demonstrated robust and comparable individual-level asymmetry, indicating that sex does not significantly influence the strength of paw preference.

#### 3.9 B. Population-Level Asymmetry

Figures 14 –17 compare right- and left-paw preference between male and female rats across all cohorts (n = 2,288). Males exhibited right-paw preference in 47% (95% CI: 43–54%) and left-paw preference in 40% (95% CI: 35–45%), while females showed comparable rates of right-paw (48%) and left-paw (39%) preference. Subgroup analyses revealed no significant sex differences for right-paw preference (χ² = 0.03, df = 1, p = 0.56 for males; χ² = 1.53, df = 1, p = 0.22 for females; Figures 14 and 16) or for left-paw preference (χ² = 0.22, df = 1, p = 0.64 for males; χ² = 0.64, df = 1, p = 0.42 for females; Figures 15 and 17). These findings indicate that sex does not systematically influence population-level asymmetry, with both males and females exhibiting comparable distributions of right- and left-paw preference.

### 3.10 Age Effects

### 3.10 A. Individual-Level Asymmetry (Young and Adult Rats)

Figures 18 and 19 summarize (n = 1,689 rats) assessing consistent paw preference in young and adult rats. Young rats exhibited a pooled prevalence of 84% (95% CI: 77–90%), while adult rats showed a comparable prevalence of 84% (95% CI: 75–90%). Heterogeneity was moderate across subgroups (I² = 40.2%–59.4%), and the subgroup difference was not significant (χ² = 0.18, df = 1, p = 0.67). These findings indicate that age does not significantly influence the strength of individual-level lateralization, with both young and adult rats showing robust and stable paw preference.

#### 3.10 B. Population-Level Asymmetry by Age

Figures 20–23 compare right- and left-paw preference across age groups. Right-paw preference was observed in 47% of rats (95% CI: 42–53%), with substantial heterogeneity (I² = 82.6%). Subgroup analyses revealed no significant difference between young (45%) and older rats (48%) (χ² = 0.41, df = 1, p = 0.52; Figure 20). Left-paw preference showed similar stability, with a pooled prevalence of 40% (95% CI: 35–45%) and no significant subgroup difference between young and older rats (χ² = 2.00, df = 1, p = 0.16; Figure 21). Comparisons between adult and other age groups also revealed no significant differences for either right- (χ² = 0.01, df = 1, p = 0.91; Figure 22) or left-paw (χ² = 2.00, df = 1, p = 0.16; Figure 23) preference. Overall, these findings indicate that paw preference remains developmentally stable once established, with age exerting minimal influence on the strength or direction of asymmetry.

### 3.11 Summary of Subgroup Analyses

Supplementary Tables 1 summarize individual and population-level findings. Table S1 presents individual-level asymmetries, showing that most rats exhibit consistent paw preference (∼84%), with subgroup differences by strain, test paradigm, sex, and age largely non-significant but with varying levels of heterogeneity. Table S2 summarizes population-level right and left paw prevalence across the same subgroups. Across test paradigms, strains, sexes, and ages, pooled estimates clustered around 45–50% for right paw and ∼39–40% for left paw, indicating no systematic directional bias at the population level. High heterogeneity (I²) across many comparisons highlights substantial methodological and strain-driven variability. Together, these results reinforce the reliability of individual-level asymmetry while underscoring the absence of consistent population-wide lateralization.

### 3.12 Sensitivity analysis

Sensitivity analyses were conducted to test the robustness of pooled prevalence estimates by systematically excluding studies identified as high risk of bias in the SYRCLE assessment and by omitting small-sample studies (n < 10). The exclusion of high-risk studies (n = 6) did not materially alter the overall results. The pooled prevalence of individual-level paw preference remained at 83% (95% CI: 77–89%, p < 0.0001) compared to 84% (95% CI: 78–89%) in the full dataset. Likewise, pooled estimates for right- and left-pawedness at the population level changed minimally (right paw: 48% → 47%; left paw: 39% → 38%), with heterogeneity indices (I²) remaining within ±3% of the original analyses. Further sensitivity testing using leave-one-out analysis confirmed that no single study exerted a disproportionate influence on pooled results; the pooled effect size fluctuated by less than ±2% across iterations.

## 4. Discussion

### 4.1 Individual and Population-Level Asymmetry

This meta-analysis provides the first comprehensive synthesis of nearly a century of rat paw-preference research, clarifying the nature and scope of hemispheric lateralization in this species. Across 40 studies encompassing 1,609 rats, we found that approximately 84% of individuals exhibited a consistent paw preference, confirming robust individual-level lateralization. These findings align with previous literature demonstrating robust individual level asymmetries.^16^ However, pooled population-level estimates revealed no significant rightward or leftward bias (right paw: 48%, left paw: 39%), with wide prediction intervals and substantial between-study heterogeneity (I² > 80%). These findings establish that while lateralized motor behavior is a conserved trait in rats, population-level handedness so prominent in humans is not a reproducible characteristic in this species. This dissociation between individual and group-level asymmetry underscores a key species difference in how motor lateralization manifests behaviorally and evolutionarily.

The absence of a population-wide directional bias in rats aligns with evolutionary evidence suggesting that strong species-level asymmetries are exceptional, largely restricted to humans and selected primates.^2, 11^ In most vertebrates, lateralization emerges as an individual trait rather than a population norm, reflecting adaptive diversification of neural control rather than evolutionary convergence on one dominant hemisphere.^3, 5, 17^ From an evolutionary standpoint, population-level handedness may have arisen in species facing high demands for social coordination or tool use; pressures largely absent in rodents.^7, 90–92^ Thus, the rat’s balanced distribution of right and left pawedness likely reflects functional lateralization without social lateralization, supporting its utility for modeling individual hemispheric specialization rather than collective motor bias.

Consistent individual-level lateralization likely stems from asymmetric organization of cortico-striatal and nigrostriatal circuits. Paw preference has been correlated with hemispheric differences in dopaminergic activity, particularly within the striatum and substantia nigra, and with asymmetrical cortical connectivity influencing skilled motor output.^26, 27, 83^ Rats displaying strong pawedness show lateralized dopamine turnover and motor-cortex dendritic morphology. Reinforcing behavioral asymmetry reflects underlying neurochemical and structural specialization.^86, 89^ In this context, the stability of individual preference observed here validates the use of pawedness as a behavioral index of hemispheric dominance, bridging cortical and subcortical lateralization mechanisms. These neurobiological substrates also underscore why rats, despite lacking population-level handedness, remains a powerful model for investigating hemispheric specialization and its dysregulation in disease.

### 4.2 Strain and Methodological Effects

A primary contributor to inter-study variability was the interaction between strain background and behavioral test paradigm. Sprague Dawley rats generally displayed more balanced paw use, whereas non–Sprague Dawley strains such as Wistar and Long–Evans exhibited mild but consistent rightward tendencies. Although these subgroup differences did not reach statistical significance, they likely reflect strain-dependent variation in dopaminergic asymmetry, stress responsivity, and learning style, as previously documented in rodent neurobehavioral studies.^16, 93^

These findings underscore that genetic background and breeding history subtly modulate the expression of lateralized behavior, warranting explicit consideration of strain as a biological variable in experimental design.

Among available rat strains, Sprague Dawley remains the most widely used in behavioral neuroscience due to its docile temperament, moderate activity, and low stress reactivity.^94^ In contrast, Wistar, Long–Evans, and Lister Hooded rats differ in exploratory behavior, anxiety profiles, and dopaminergic activity, all these factors can influence laterality strength and consistency.^95–99^ Also, Sprague Dawley rats dominate toxicological and parkinsonian modeling studies. Establishing their baseline lateralization profile is critical for interpreting hemispheric lesion, toxin, or transplantation outcomes.^39, 100–102^ Without this contextual baseline, inter-strain variability can be mistaken for treatment effects, leading to spurious conclusions about hemispheric dominance.

Similarly, from a methodological perspective, test paradigm selection further amplified heterogeneity. The Collins food-reaching test, historically the most standardized and widely adopted assay, was chosen for cross-study comparison because of its simplicity, repeatability, and quantifiable reach count, allowing direct comparability across decades of research.^61, 63–65, 69^ However, this test often produced slightly weaker asymmetries compared with skilled-reaching paradigms such as the Staircase test, Pawedness Trait test, and Vermicelli Handling task, which demand higher sensorimotor precision and repetitive forelimb coordination. These complex tasks recruit broader cortico-striatal circuitry and proprioceptive feedback mechanisms, thereby enhancing sensitivity to detect lateralized motor control.

Collectively, these findings indicate that both strain background and task complexity interact to shape the magnitude but not the direction of paw preference. Future research should adopt standardized, multi-paradigm behavioral batteries (e.g., combining Collins and skilled-reaching tests) while ensuring transparent reporting of strain, age, and sex. Such standardization will minimize methodological bias, enhance cross-study comparability, and yield a more reliable understanding of hemispheric asymmetry in rodent models.

### 4.3 Sex and Age Effects

Sex and age exerted subtle but biologically meaningful effects. Female rats showed marginally greater consistency in paw preference (92%) than males (83%), with lower heterogeneity across studies, consistent with prior findings suggesting sex hormone modulation of motor lateralization.^103^ Estrogen has been implicated in enhancing hemispheric asymmetry in motor and cognitive tasks, potentially influencing paw-use stability.^71^ Conversely, age effects were negligible: both young and adult rats exhibited comparable prevalence of paw preference (∼84%), suggesting that once lateralization is established, it remains developmentally stable. This aligns with longitudinal behavioral evidence indicating that motor laterality in rodents consolidates early and persists across the lifespan.^3^ The limited availability of balanced male–female and age-stratified cohorts, however, remains a key gap in the literature, warranting systematic control in future behavioral studies.

### 4.4 Translational Implications

The translational implications of these findings extend beyond behavioral neuroscience. In disorders characterized by asymmetric onset, most notably Parkinson’s disease (PD), where dopaminergic degeneration often begins unilaterally understanding species-specific lateralization is critical for model fidelity.^37, 38, 86, 104^ Our synthesis supports the rat as a reliable model for individual hemispheric specialization but not for population-level handedness. This distinction carries practical implications: preclinical PD models relying on unilateral 6-hydroxydopamine (6-OHDA) lesions or paraquat–lectin combinations may better approximate early-stage asymmetry when strain and task selection account for intrinsic lateralization tendencies.^102, 105^

Moreover, variability across strains and test paradigms in this meta-analysis mirrors the clinical heterogeneity of hemispheric vulnerability, emphasizing the need to align behavioral laterality measures with neuroanatomical correlates of dopaminergic asymmetry. Such as striatal tyrosine hydroxylase (TH), vesicular monoamine transporter 2 (VMAT2), and dopa decarboxylase (DDC) expression. Notably, rats with dominant right paw use have been shown to exhibit higher left striatal TH expression, supporting a direct behavioral–molecular correspondence that strengthens the translational bridge between lateralized motor behavior and hemispheric dopaminergic function.^26^

### 4.5 Limitations and Future Directions

Risk of bias assessments revealed frequent gaps in the reporting of randomization, blinding, and allocation concealment, which may contribute to variability and reduce confidence in the pooled findings. These methodological inconsistencies, together with substantial between-study heterogeneity, make direct comparisons across experiments difficult. Other limitations include the reliance on imputed data when raw counts were not available, underpowered subgroup analyses due to uneven representation of strain, sex, and age groups, and the inability to perform meta-regression because of incomplete reporting. Variation in behavioral scoring criteria, particularly in older studies, also contributes to differences in effect estimates.

Improving reproducibility in future paw preference research will require greater standardization of testing procedures, such as combining the Collins test with skilled-reaching paradigms, as well as consistent reporting of raw paw-use counts and balanced cohort design across strains and sexes. Clear documentation of randomization and blinding practices might strengthen methodological rigor. Although this review covered studies spanning nearly a century, limited reporting of variance prevented formal assessment of publication bias using funnel plots or Egger’s regression. Since paw preference is often reported as a secondary outcome, selective omission cannot be excluded. However, the presence of studies showing rightward, leftward, and balanced outcomes, along with sensitivity analyses showing stable pooled estimates after excluding smaller or higher risk-of-bias studies, suggests that the central conclusions are robust. Future studies that routinely report raw counts and variance will allow more definitive evaluation of publication bias in subsequent meta-analyses.

## 5. Conclusion

This systematic review and meta-analysis demonstrate that rats consistently exhibit strong individual-level paw preference, confirming their validity as a behavioral marker of hemispheric lateralization. However, no consistent population-wide right- or left-paw bias was observed, underscoring a fundamental species difference compared to the pronounced right-handedness in humans. Subgroup analyses revealed modest influences of strain and test paradigm, while sex-and age-related effects were subtle and inconsistent. From a translation perspective, paw preference remains a valuable tool for investigating hemispheric specialization and asymmetric disease onset. Yet the absence of universal population-level asymmetry highlights the need for careful interpretation when extrapolating rodent findings to human contexts. To maximize reproducibility and translational value, future studies should adopt standardized behavioral protocols, report raw paw counts, include balanced cohorts across strains and sexes, and improve transparency in randomization and blinding. These steps will reduce heterogeneity, strengthen cross-study comparisons, and enhance the reliability of paw preference as a model for lateralization in neuroscience research.

## Abbreviations

CI confidence interval
PRISMA Preferred Reporting Items for Systematic Reviews and Meta-Analyses
LI laterality index
CAS coefficient of asymmetry

## Ethics statement

We confirm that we have read the Journal’s Position on Ethics in Publishing and that this manuscript is consistent with those guidelines.

## CRediT authorship contribution statement

Conceptualization : DP, TS ; Data Curation : DP, CSS ; Formal Analysis : DP and CCS ; Funding acquisition : TS ; Investigation : DP, CCS, DB, MRR ; Methodology : DP, CSS ; Project administration : DP ; Resources : TS ; Supervision : DP and TS ; Visualization : DP ; Writing of original draft : DP ; Reviewing and Editing : DP and TS.

## Declaration of generative AI use

During the preparation of this work, the authors used Perplexity AI to assist with grammar and language editing. Following its use, the authors reviewed and revised the content as necessary and took full responsibility for the integrity and accuracy of the final manuscript.

## Data availability

The data that supports the findings of this study are available on request from the corresponding author.

## Acknowledgements

Dr. Sadik A. Khuder, PhD, Professor of Medicine, Statistics, and Public Health at the University of Toledo College of Medicine and Life Sciences, 43614. for his assistance with statistical methodology. Dr. Rebecca L. Morgan PhD, MPH, Professor of Population and Quantitative Health Sciences, School of Medicine, Case Western Reserve University, 44106. for her guidance in early planning and methodology.

