## Supplementary_Table_1 for "Paw Preference in Rats Across Tests, Strains, Sex, and Age: A PRISMA-Compliant Systematic Review and Meta-Analysis"

Summary of Subgroup Analyses:

**Table: S1. Individual-Level Subgroup Analyses**

| **Comparison** | **Subgroup** | **Pooled Proportion** | **95% CI** | **Heterogeneity (I²)** |
| --- | --- | --- | --- | --- |
| Test Paradigm | Collins | 85% | 80–89% | 76.1% |
| Test Paradigm | Other tests | 82% | 56–94% | 93.9% |
| Strain | Sprague Dawley | 76% | 54–89% | 95.6% |
| Strain | Other strains | 87% | 81–91% | 78.1% |
| Sex | Male | 83% | 77–89% | 63.8% |
| Sex | Female | 92% | 85–95% | 9.5% |
| Age | Young | 84% | 77–90% | 40.2% |
| Age | Adult | 84% | 75–90% | 59.4% |

Note: I² values represent heterogeneity across included studies. Thresholds: 25% = low, 50% = moderate, 75% = high heterogeneity.

**Table: S2. Population-Level Subgroup Analyses**

| **Comparison** | **Subgroup** | **Pooled Prevalence** | **95% CI** | **Heterogeneity (I²)** |
| --- | --- | --- | --- | --- |
| Right Paw (Test Paradigm) | Collins | 48% | 41–54% | 84.1% |
| Right Paw (Test Paradigm) | Other tests | 49% | 38–60% | 78.6% |
| Left Paw (Test Paradigm) | Collins | 41% | 35–47% | 80.5% |
| Left Paw (Test Paradigm) | Other tests | 34% | 23–45% | 69.1% |
| Right Paw (Strain) | Sprague Dawley | 45% | 33–58% | 85.7% |
| Right Paw (Strain) | Other strains | 49% | 43–54% | 81.8% |
| Left Paw (Strain) | Sprague Dawley | 35% | 26–45% | 74.6% |
| Left Paw (Strain) | Other strains | 40% | 35–44% | 77.1% |
| Right Paw (Sex) | Male | 47% | 40–54% | 76.6% |
| Right Paw (Sex) | Other | 45% | 41–50% | 87.8% |
| Left Paw (Sex) | Male | 40% | 32–49% | 85.4% |
| Left Paw (Sex) | Other | 38% | 33–44% | 72.6% |
| Right Paw (Sex) | Female | 60% | 42–77% | 83.6% |
| Right Paw (Sex) | Other | 47% | 41–52% | 81.6% |
| Left Paw (Sex) | Female | 30% | 16–51% | 84.6% |
| Left Paw (Sex) | Other | 40% | 35–44% | 77.3% |
| Right Paw (Age) | Young | 45% | 33–57% | 71.3% |
| Right Paw (Age) | Other | 48% | 42–55% | 82.6% |
| Left Paw (Age) | Young | 39% | 30–49% | 73.3% |
| Left Paw (Age) | Other | 39% | 35–44% | 79.9% |
| Right Paw (Age) | Adult | 49% | 43–54% | 84.2% |
| Right Paw (Age) | Other | 45% | 37–53% | 71.3% |
| Left Paw (Age) | Adult | 39% | 34–44% | 79.9% |
| Left Paw (Age) | Other | 40% | 31–50% | 73.3% |

Note: Population-level analyses reveal balanced right vs. left paw prevalence across subgroups, with no consistent directional bias. High I² values highlight methodological and strain-driven variability.
